# ANKRD31 regulates spatiotemporal patterning of meiotic recombination initiation and ensures recombination between heterologous sex chromosomes in mice

**DOI:** 10.1101/423293

**Authors:** Frantzeskos Papanikos, Julie A.J. Clément, Erika Testa, Ramya Ravindranathan, Corinne Grey, Ihsan Dereli, Anastasiia Bondarieva, Sarai Valerio-Cabrera, Marcello Stanzione, Alexander Schleiffer, Petr Jansa, Diana Lustyk, Fei Jifeng, Jiri Forejt, Marco Barchi, Bernard de Massy, Attila Toth

**Author notes:** These authors contributed equally to this work (Frantzeskos Papanikos, Julie A.J. Clément).

## Abstract

Orderly segregation of chromosomes during meiosis requires that crossovers form between homologous chromosomes by recombination. Programmed DNA double-strand breaks (DSBs) initiate meiotic recombination. We identify ANKRD31 as a critical component of complexes of DSB-promoting proteins which assemble on meiotic chromosome axes. Genome-wide, ANKRD31 deficiency causes delayed recombination initiation. In addition, loss of ANKRD31 alters DSB distribution owing to reduced selectivity for sites that normally attract DSBs. Strikingly, ANKRD31 deficiency also abolishes uniquely high rates of recombination that normally characterize pseudoautosomal regions (PARs) of X and Y chromosomes. Consequently, sex chromosomes do not form crossovers leading to chromosome segregation failure in ANKRD31-deficient spermatocytes. These defects are accompanied by a genome-wide delay in assembling DSB-promoting proteins on axes and a loss of a specialized PAR-axis domain that is highly enriched for DSB-promoting proteins. Thus, we propose a model for spatiotemporal patterning of recombination by ANKRD31-dependent control of axis-associated complexes of DSB-promoting proteins.

## Highlights

Temporal and spatial patterning of recombination are regulated by ANKRD31

Selective use of PRDM9 binding sites as DSB hotspots requires ANKRD31

Enrichment of pro-DSB factors in the PAR requires ANKRD31 but not IHO1

Recombination in the PAR critically depends on ANKRD31

## Introduction

Programmed DNA double-strand breaks (DSBs) are critical for meiosis, as they initiate meiotic homologous recombination, and their repair generates reciprocal DNA exchanges, called crossovers (CO). In most organisms, at least one CO must form between each pair of homologous chromosomes (homologs) for correct segregation of chromosomes in the first meiotic division (Gray and Cohen, 2016; Hunter, 2015). Persistent DSBs are potentially genotoxic; anomalies in CO formation cause aneuploidies and infertility. Hence, meiotic DSB formation and repair are under tight spatiotemporal control (Keeney et al., 2014).

Meiotic DSB formation and repair take place within the context of the meiosis-specific chromosome axis and the synaptonemal complex (SC), reviewed in (de Massy, 2013; Hunter, 2015; Keeney et al., 2014). Initially, multiple DSBs are formed along the axes, which are linear chromatin structures that assemble along the shared core of sister chromatid pairs early in meiotic prophase. Single-stranded DNA ends formed by processing of DSBs invade homologs, which promotes alignment and synapsis of homolog axes. It is the resultant SC where recombination completion, including CO formation, occurs.

DSB formation is catalyzed by an evolutionary conserved topoisomerase-like enzyme complex, consisting of the SPO11 enzyme and its binding partner TOPOVIBL (Baudat et al., 2000; Bergerat et al., 1997; Keeney et al., 1997; Robert et al., 2016; Romanienko and Camerini-Otero, 2000; Vrielynck et al., 2016). SPO11 activity depends on a partially conserved set of auxiliary proteins (de Massy, 2013). Before DSB formation starts, axes recruit pre-DSB recombinosomes, which are complexes of three conserved and essential DSB-promoting proteins, MEI4, REC114 and IHO1 in mice (de Massy, 2013; Kumar et al., 2010; Kumar et al., 2018; Stanzione et al., 2016). These pre-DSB recombinosomes are thought to promote DSBs by activating SPO11 on axes. Anchoring of pre-DSB recombinosomes to axes relies on IHO1 (Mer2 in budding yeast) and meiosis-specific HORMA-domain proteins in both budding yeast and mammals (Kumar et al., 2015; Panizza et al., 2011; Stanzione et al., 2016). HORMAD1-IHO1 assemblies can form independent of MEI4 or REC114 thereby providing an axis-bound platform for pre-DSB recombinosome assembly in mice (Kumar et al., 2018; Stanzione et al., 2016). HORMAD1, IHO1 and pre-DSB recombinosomes are restricted to unsynapsed axes, which is hypothesized to concentrate DSB forming activity to genomic regions requiring DSBs for completion of homolog alignment (Kauppi et al., 2013; Stanzione et al., 2016; Wojtasz et al., 2009).

Meiotic chromatin is thought to be arrayed into loops anchored on chromosome axes (Kleckner, 2006; Zickler and Kleckner, 1999) and DSBs seem to form in these loops (Blat et al., 2002; Panizza et al., 2011). Sites of frequent DSB formation, called hotspots, are also associated with open chromatin marks, such as trimethylation of histone H3 lysine 4 (H3K4me3) and lysine 36 (H3K36me3) (Buard et al., 2009; Diagouraga et al., 2018; Grey et al., 2011; Powers et al., 2016; Smagulova et al., 2011). Thus, axis-bound pre-DSB recombinosomes are hypothesized to target SPO11 activity by recruiting open chromatin sites from within loops to the axis via a poorly understood mechanism, reviewed in (Grey et al., 2018). In most mammals, DSB hotspots emerge at sites bound by the histone methyltransferase PRDM9 in a sequence specific manner. PRDM9 generates the histone marks H3K4me3 and H3K36me3 at these sites (Grey et al., 2018). Additionally, by an unknown mechanism, PRDM9 prevents DSBs at other H3K4me3-rich genomic locations such as transcription start sites or enhancers, which serve as hotspots in *Prdm9^-/-^* meiocytes (hereafter called default hotspots) (Brick et al., 2012; Diagouraga et al., 2018). An exception to this rule is the pseudoautosomal region (PAR) of X and Y chromosomes, where high DSB activity occurs independently of PRDM9 (Brick et al., 2012). X and Y chromosomes are homologous only in the PAR, therefore at least one CO must form in this relatively short region to ensure faithful segregation of these chromosomes. Thus, DSB density is between 10 to 110 fold higher in the PAR than the rest of the genome (Kauppi et al., 2011; Lange et al., 2016). High DSB activity is possibly enabled by a unique PAR-chromatin environment characterized by both strong accumulation of PRDM9-independent histone H3K4me3 (Brick et al., 2012), and the combination of a disproportionately long axis and short chromatin loops (Kauppi et al., 2011). However, the importance of these features in DSB control has not been tested, and mechanisms underlying high recombination rate in the PAR remain enigmatic.

Here we report the identification of a novel meiotic protein, ANKRD31, which plays a major role in the temporal and spatial control of meiotic DSBs. Strikingly, ANKRD31 is required to prevent DSBs at default hotspot sites, and to ensure DSBs/COs in the PAR. Both roles likely rely on ANKRD31’s function in organizing axis-bound pre-DSB recombinosomes.

## Results

### Conserved ANKRD31 associates with MEI4 and REC114-marked protein complexes along unsynapsed chromosome axes

To identify new players in meiotic recombination we searched for genes that are preferentially expressed in mouse gonads where prophase stage meiocytes are present (Finsterbusch et al., 2016; Papanikos et al., 2016; Wojtasz et al., 2009). We reviewed ENCODE transcriptome datasets (Consortium, 2012; Sloan et al., 2016) and carried out expression profiling of reproductive tissues. One of the genes identified by this analysis was *Ankrd31* (Figure 1A, 1B and 1SA). The largest open reading frame (ORF) of *Ankrd31* encodes a 1857 amino acid protein that is conserved among vertebrates. A multiple alignment of ANKRD31 sequences identified five conserved regions (Figure 2S). In most vertebrate taxa, ANKRD31 contains two separated triplets of Ankyrin repeats, which are common protein-protein interaction mediator motifs (Mosavi et al., 2004). C-terminal to the second Ankyrin repeat domain there is a predicted coiled-coil domain and two further regions, CR4 and CR5, without functional predictions.

**Figure 1.**
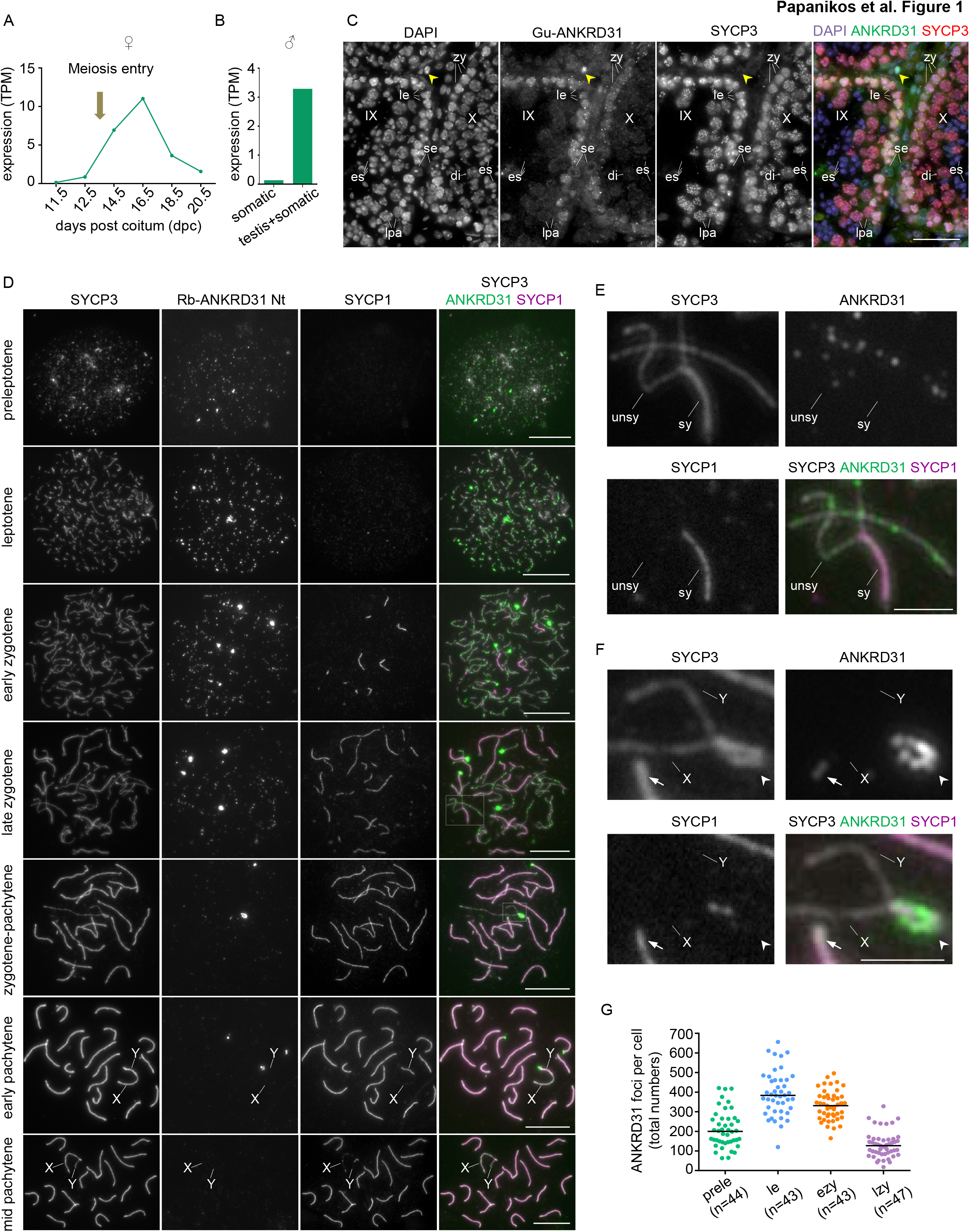
ANKRD31 is a meiotic protein that preferentially localizes to unsynapsed chromosome axes. **(A-B)** *Ankrd31* transcript levels are shown as transcripts per million (TPM) in (**A**) total RNAs of fetal ovaries at the indicated developmental stages, or (**B**) mixtures of equal quantities of total RNAs of 17 somatic tissues (somatic) or 17 somatic tissues plus adult testes (testis+ somatic). (**C**) DAPI detected DNA on cross sections of adult mouse testes. Immunofluorescence detected ANKRD31 and a marker of meiotic chromosome axis, SYCP3. Antibodies raised against an N-terminal fragment of ANKRD31 in guinea pig (Gu-ANKRD31 Nt) were used. Seminiferous tubules at stage IX and X are shown; see STAR Methods for seminiferous tubule staging. Sertoli cells (se), leptotene (le), zygotene (zy), late pachytene (lpa), and diplotene (di) spermatocytes and elongating spermatids (es) are marked. A strong signal in some interstitial cells (yellow arrowhead) and a weak nuclear signal in the nucleus of Sertoli cells were detected when sections were stained by Gu-ANKRD31 Nt antibodies. We consider these signals non-specific as rabbit antibodies raised against the same fragment of ANKRD31 (Rb-ANKRD31 Nt) did not produce this pattern, see Figure S1B. (**D-F**) Indicated proteins were detected in nuclear surface spread spermatocytes of adult mice at the indicated prophase stages. SYCP1 is a marker of synapsed chromosomal regions. Rb-ANKRD31 Nt antibodies detected ANKRD31. Matched exposed images are shown for ANKRD31. SYCP3 and SYCP1 were differentially levelled to optimize simultaneous observation of SYCP3, SYCP1 and ANKRD31 in overlay images. Insets of late zygotene and zygotene-pachytene stage spermatocytes are enlarged in **E** and **F**, respectively. X and Y chromosomes and unsynapsed (unsy) and synapsed (sy) regions of an autosome are indicated in some images of **D-F**. In **F**, arrowhead and arrow mark a large and a small aggregate of ANKRD31 on the PAR ends of sex chromosomes and the end of an autosome, respectively. Bars, 50μm (**C**), 10μm (**D**), 2.5μm (**E, F**). (**G**) Quantification of total ANKRD31 focus numbers (Rb-ANKRD31 Ct) in nuclear surface spread preleptotene (prele), leptotene (le), early zygotene (ezy) and late zygotene (lzy) spermatocytes. Number of cells and medians are indicated. See also Figure S1, S2 and Table S1.

To test if ANKRD31 functions in meiotic cells we stained ANKRD31 in histological sections of adult mouse testes with antibodies raised to N- or C-terminal fragments of ANKRD31 (Figure 1C and S1B). Somatic cells apparently lacked specific ANKRD31 signal as distinct antibodies produced inconsistent staining patterns. In contrast, antibodies to both an N- and a C-terminal fragment of ANKRD31 produced consistent staining in spermatocyte nuclei from premeiotic DNA replication until early pachytene. Localization of ANKRD31 was further analyzed in nuclear surface spreads of spermatocytes where detection of the chromosome axes and SC allow meiotic prophase to be sub-staged (Figure 1D and Table S1, see also STAR Methods for staging prophase). ANKRD31 was detected on chromatin between preleptotene and early pachytene in spermatocyte spreads by both the anti-N-terminal and anti-C-terminal ANKRD31 antibodies (Figure 1D-G and S1C-E). ANKRD31 focus numbers gradually increased from preleptotene until leptotene and early zygotene where median ANKRD31 foci numbers peaked (Figure 1D and 1G). The majority of foci associated with the assembling chromosome axes from leptotene onwards (85.5% and 88.7% in leptotene, n=29, and early zygotene, n=32, spermatocytes, respectively). ANKRD31 foci were depleted from synapsed axes (Figure 1E, S1C and S1D), and ANKRD31 focus numbers dropped as cells progressed in zygotene and beyond, which resembles the behaviour of pre-DSB recombinosomes (Kumar et al., 2010; Kumar et al., 2015; Stanzione et al., 2016). We found very substantial colocalization between ANKRD31 and pre-DSB recombinosomes detected by MEI4 or REC114 staining in leptotene and early zygotene spermatocytes (Figure 2A-F). ANKRD31 also formed foci in pre-pachytene stage oocytes on unsynapsed axes, and these foci also colocalized with MEI4 (Figure S3A). Thus, ANKRD31 associates with pre-DSB recombinosomes in both sexes.

**Figure 2.**
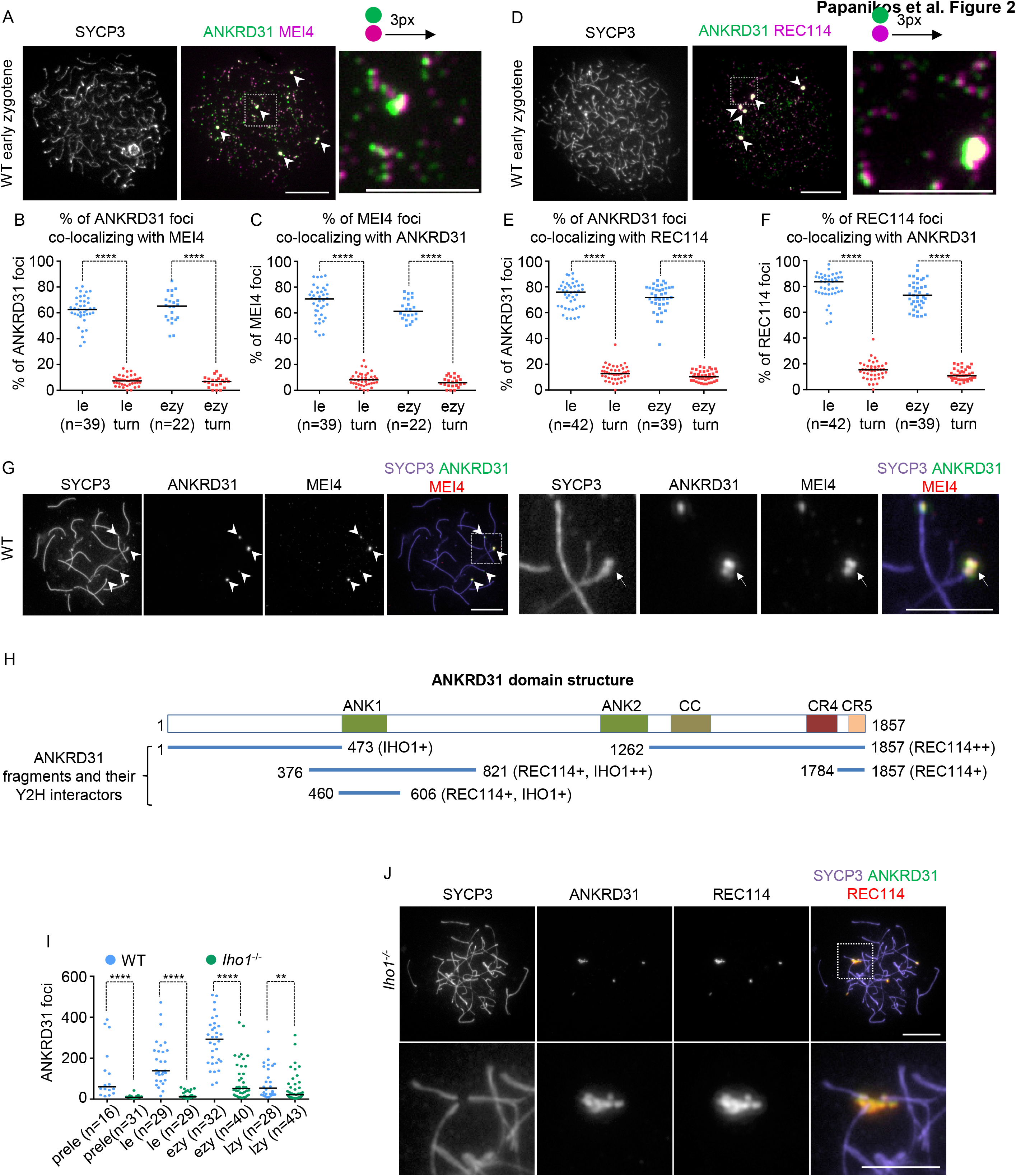
ANKRD31 colocalizes with DSB-promoting proteins on chromosome axes, and it forms large aggregates at PARs and PAR-like genomic regions. (**A, D, G, J**) Indicated proteins were detected by immunofluorescence in nuclear spread spermatocytes. MEI4 (**A**) and REC114 (**D**) signal is shifted to the right by three pixels in the enlarged insets. Arrowheads mark large co-aggregates of ANKRD31, MEI4 and REC114. (**B, C, E, F**) Quantification of overlap between ANKRD31 foci and either MEI4 (**B, C**) or REC114 (**E, F**) foci in leptotene (le) and early zygotene (ezy). Colocalization was significantly reduced when ANKRD31 images were turned 90 degree (turn, red data points) relative to MEI4 or REC114 images. Four stars indicate P<0.0001, Wilcoxon matched-pairs signed rank test. (**G**) Coaggregates of ANKRD31 and MEI4 are shown (marked by arrowheads) at the ends of three autosomes and the paired PARs of sex chromosomes in a spermatocyte at zygotene-to-pachytene transition. The four images on the right show an enlarged version of boxed area in the left panel and arrow marks PAR. (**H**) Schematic representation of ANKRD31 domain structure. Colored boxes mark the first (ANK1) and second (ANK2) sets of Ankyrin repeats, a conserved coiled coil domain (CC), and conserved regions four (CR4) and five (CR5). Blue lines and numbers indicate ANKRD31 protein residues that interact with the indicated proteins in yeast two-hybrid (Y2H). + and ++ mark weak and strong interactions, respectively. (**I**) Focus counts of ANKRD31 in *Iho1^+/++^* (WT, blue) and *Iho1^-/-^* (green) spermatocytes in preleptotene (prele), leptotene (le), early (ezy) and late (lzy) zygotene. Counted number of cells and medians are indicated. ** and *** indicate significance at 0.001<P<0.05 and P<0.0001, Mann-Whitney U test. (**J**) *Iho1^-/-^* spermatocyte with fully formed chromosome axes. Enlarged insets (bottom panel) show a co-aggregate of ANKRD31 and REC114. Bars are 10μm and 5μm in low resolution images and enlarged insets in **A**, **D**, **G** and **J**. See also Figure S3, Table S2 and S3.

We used yeast two-hybrid (Y2H) assays (Figure 2H, S3D and Table S2) to test if ANKRD31 could directly interact with known components of pre-DSB recombinosomes, or with HORMAD1, which acts as an anchor for pre-DSB recombinosomes on axes (Kumar et al., 2015; Stanzione et al., 2016). We found that fragments of ANKRD31 that contained the N-terminal Ankyrin repeats interacted with both IHO1 and REC114. REC114 also interacted with the CR5 domain of ANKRD31. These results suggest that ANKRD31 might be incorporated into pre-DSB recombinosomes by directly interacting with REC114 and IHO1. This conclusion is also supported by similar and complementing observations of the accompanying manuscript of Boekhout et al.

### ANKRD31 aggregates form at PARs

Beyond numerous relatively small foci (median diameter, 0.26 μm), ANKRD31 also formed relatively large aggregates (median diameter 0.62 μm) in smaller numbers (up to 8), which were detectable from preleptotene to pachytene (Figure 1D, 1F and S1E). These large ANKRD31 aggregates always colocalized with similar aggregates of REC114 (n=98 spermatocytes) and MEI4 (n=89 spermatocytes and n=24 oocytes, Figure 2A, 2D, 2G and 3SA-C), and declined in median number from five to three as spermatocytes progressed from leptotene to early pachytene (n=108 spermatocytes). In zygotene and pachytene, where chromosome axes were well developed, it was apparent that these ANKRD31 aggregates associated with the ends of a subset of chromosomes (Figure 1D and 1F). One of the ANKRD31 aggregates always associated with the PARs on the X and Y chromosomes in spermatocytes (Figure 1D, 1F, 2G and S3C), and ANKRD31 aggregates were also associated with up to three non-centromeric autosomal ends (Figure S1E). Peculiarly, whereas the small ANKRD31 foci were detectable only on unsynapsed axes, ANKRD31, MEI4, and REC114 aggregates persisted on synapsed autosomal ends and PARs as meiocytes progressed to early pachytene (Figure 1D, 1F, 2G and S3B-C). This was most obvious in the PARs of spermatocytes (see high magnification images in Figure 1F, 2G and S3C). These large aggregates of ANKRD31, MEI4 and REC114 disappeared as meiocytes progressed beyond early pachytene (Figure 1D and not shown). Our observations are consistent with unpublished data of Acquaviva, Jasin & Keeney (personal communication). Acquaviva et al. observed ANKRD31 aggregates on PARs, and identified chromosome 4, 9 and 13 as autosomes whose non-centromeric ends carry arrays of PAR-like sequences that associate with ANKRD31 aggregates.

### Distinct molecular requirements for ANKRD31 aggregates and ANKRD31 foci

Given their distinct behaviour in synapsed regions, foci and aggregates of ANKRD31/MEI4/REC114 might represent qualitatively different protein complexes with distinct underlying molecular requirements. To test this possibility we compared localization of ANKRD31, REC114 and MEI4 in *Mei4^-/-^, Rec114^-/-^* and *Iho1^-/-^* spermatocytes (Figure 2J and S3E-I). The numbers of REC114 and MEI4 foci are strongly reduced in *Mei4^-/-^* and *Rec114^-/-^* mice, respectively (Kumar et al., 2018). In addition, we found that both focus and aggregate formation of ANKRD31, REC114 and MEI4 were each disrupted in *Mei4^-/-^* and *Rec114^-/-^* spermatocytes (Figure S3E-G and Table S3). Remarkably, only the formation of ANKRD31 (Figure 2I), MEI4 and REC114 foci (Stanzione et al., 2016), but not aggregates, were disrupted in *Iho1^-/-^* spermatocytes (Figure 2J). ANKRD31 aggregates formed efficiently in *Iho1^-/-^* spermatocytes; median numbers of ANKRD31 aggregates were four in both wild type (n=62) and *Iho1^-/-^* (n=57) spermatocytes in zygotene. These aggregates always colocalized with aggregates of MEI4 (n=100) and REC114 (n=100) in *Iho1^-/-^* spermatocytes (Figure 2J and S3H). ANKRD31 aggregates also colocalized with PAR FISH signals in late zygotene-like *Iho1^-/-^* spermatocytes (≥94% of FISH signals and ANKRD31 aggregates colocalize with each other, n=51) (Figure S3I). These observations support the hypothesis that ANKRD31 interacts with MEI4/REC114 complexes. Furthermore, these data show that distinct molecular requirements underpin focus formation of ANKRD31, MEI4 and REC114 in non-PAR regions and aggregate formation of the same proteins at PARs and PAR-like ends of chromosome 4, 9 and 13 (identity of these autosomes was revealed by personal communication with Acquaviva, Jasin & Keeney).

### Loss of ANKRD31 affects fertility of mice

The subcellular localization of ANKRD31 suggested that ANKRD31 was involved in meiotic recombination. To test this hypothesis we generated two mouse lines (by CRISPR/Cas9 genome editing) which carried distinct frameshift mutations in exon 6 of *Ankrd31* (Figure 3A). Initial analysis uncovered indistinguishable phenotypes between these lines, hence, we analyzed only one of them, mut1, in depth. The frame of the longest predicted ORF of the *Ankrd31* transcript, XM_006517797.1, was disrupted in the mut1 line after the 134^th^ codon (Figure 3A), which is predicted to truncate the normal protein sequence 330 amino acid upstream of the first conserved domain (Ankyrin repeat 1). We therefore refer to mut1 as a loss of function allele (*Ankrd31^-/-^*) hereafter.

**Figure 3.**
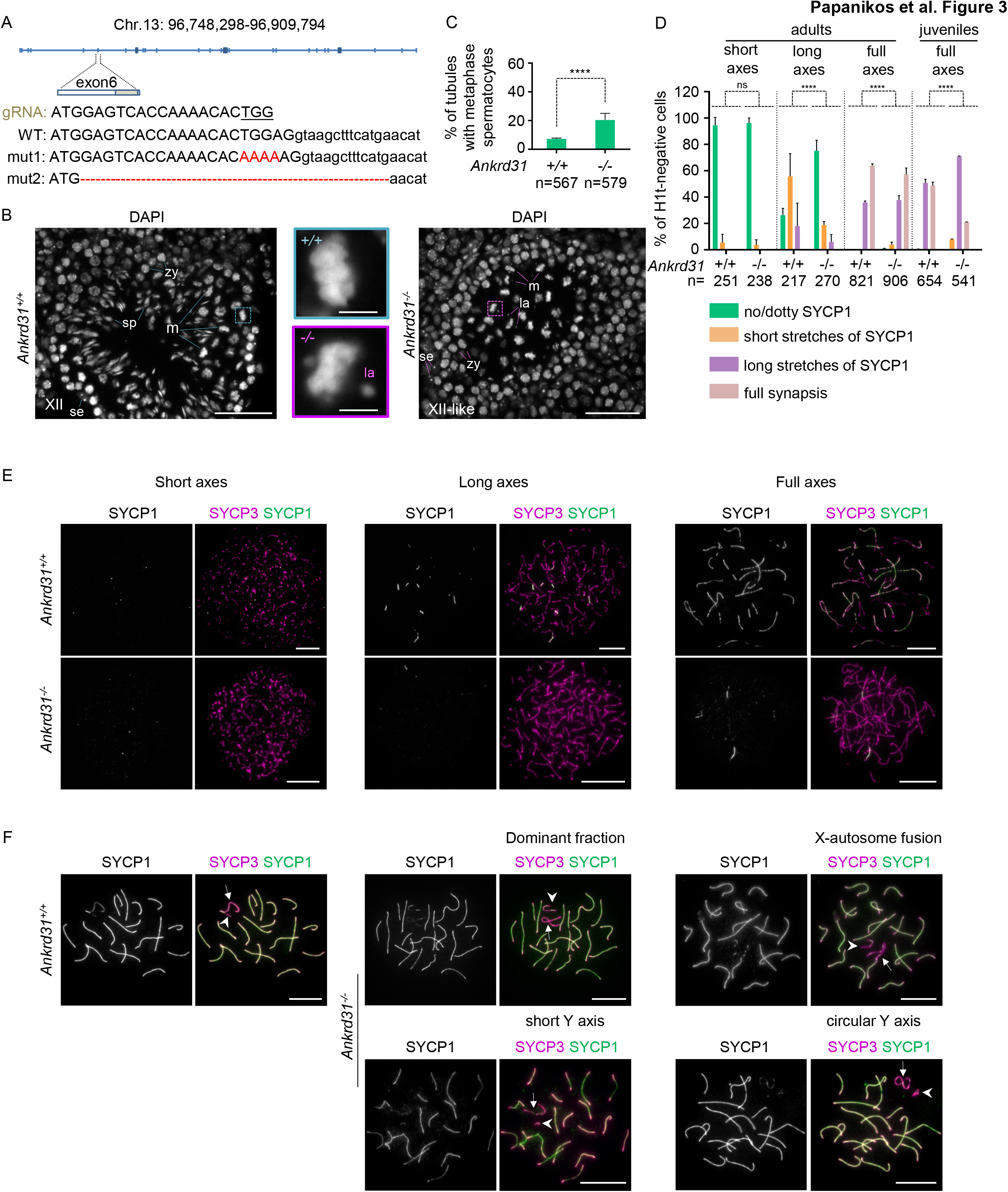
SC formation is defective in *Ankrd31^-/-^* spermatocytes. (**A**) Schematic of the *Ankrd31* locus and targeting of exon 6 by CRISPR/Cas9. Guide RNA sequence (gRNA) with PAM (underlined), the corresponding 3-prime end of exon 6 in wild-type (WT) and in two distinct *Ankrd31* mutant lines (mut1 and 2) are shown. Red marks mutated sequence. (**B**) DNA was stained by DAPI on cross sections of testes of indicated genotypes. Seminiferous tubules are shown at stages where the luminal layer of spermatocytes are in metaphase (stage XII in wild-type, XII-like in *Ankrd31^-/-^,* see STAR Methods). Sertoli cells (se), zygotene spermatocytes (zy, basal layer of spermatocytes), metaphase cells (m), lagging chromosomes (la) and immature sperm (sp) are marked. The middle panel shows enlarged images of the indicated boxed areas of the left and right panels. Bars are 50μm (sections) and 5μm (zoom) (**C**) Quantification of seminiferous tubules that contain metaphase cells in the luminal layer of spermatocytes. Number of tubules (n) counted in four experiments, the weighted averages of percentages and standard deviation is shown. (**D-F**) SC (SYCP1) was detected in combination with axis marker (SYCP3) in *Ankrd31^+/+^* and *Ankrd31^-/-^* spermatocytes. (**D**) Quantifications of SC formation in histone H1t-negative spermatocytes. SC formation was assessed by categorising SYCP1 staining patterns as indicated (full synapsis indicates SYCP1 was detected along the entire length of all autosomes). Quantification was done in subpopulations of spermatocytes that had short axis (equivalent to wild-type leptotene), long incomplete axes (equivalent to early-zygotene), or fully formed continuous axes (equivalent to late zygotene and early pachytene cells). Spermatocytes were isolated from adults or 13 day old juvenile mice as indicated. Number of counted cells (n), standard deviation and the weighted averages of three (adults) or two (juvenile, 13 days old) experiments are shown. (**E, F**) Indicated proteins are shown in nuclear spread *Ankrd31^+/+^* and *Ankrd31^-/-^* spermatocytes. (**E**) Images of spermatocytes from adults represent axis development categories that were quantified in **D**. **F** shows sex chromosome configurations of wild-type and *Ankrd31^-/-^* spermatocytes. Spermatocytes with unsynapsed X and Y chromosomes (without other apparent abnormalities) represent a dominant fraction (76%) of *Ankrd31^-/-^* spermatocytes, minor fractions are represented by spermatocytes that have extremely short or self-synapsed Y axes (12.5%), X-autosome fusions (6.6%) and apparently circularized Y (4.8%), n=271 spermatocytes. X (arrows) and Y (arrowhead) chromosomes are marked. (**E, F**) Bars, 10μm. (**C, D**) Results of Chi Square statistics are indicated as non-significant (ns) and P<0.0001 (****). See also Figure S4, S5, Table S4, and S5.

We confirmed the loss of full length ANKRD31 from *Ankrd31^-/-^* meiocytes by both immunoprecipitation-western blot analysis of testis extracts and immunofluorescence in fixed gonadal samples (Figure S4A-F). Disruption of *Ankrd31* caused no obvious somatic phenotypes but it led to fertility defects in both sexes. Whereas female *Ankrd31^-/-^* mice were fertile, they lost fertility faster than wild-type with advanced age. Average numbers of pups/breeding week were 1.53 and 1.38 between 8-28 weeks of age (not significant), and 0.92 and 0.45 beyond 28 weeks of age (P=0.0272, paired t-test) in wild-type and *Ankrd31^-/-^* mothers, respectively (n=6 mice for each genotype). Loss of ANKRD31 caused infertility in males, as no pups were observed after 34.5 breeding weeks (n=2 males).

To test if these fertility defects were caused by a reduced capacity of *Ankrd31^-/-^* meiocytes to progress through meiosis, we examined oocytes and spermatocytes in histological sections. Consistent with reduced female fertility, we observed 4.95 fold lower median oocyte numbers in *Ankrd31^-/-^* mice than wild-type at 6-7 weeks of age (Figure S4G). This reduction in oocyte number was associated with elevated rates of apoptosis in the oocytes of newborn *Ankrd31^-/-^* mice as compared to wild-type (Figure S4H). Consistent with a loss of fertility in males, testis weight was 2.7 fold lower in *Ankrd31^-/-^* than wild-type mice (Figure S4I). Investigation of gametogenesis in histological sections revealed elevated apoptosis after mid pachytene in spermatocytes of *Ankrd31^/-^* mice (Figure S4J and Table S4). Nevertheless, noticeable fractions of spermatocytes progressed to the first meiotic metaphase, where they arrested, which was evidenced by the elevated proportion of seminiferous tubules that contained metaphase cells (Figure 3B and 3C). These metaphase cells had lagging chromosomes indicating chromosome alignment defects in *Ankrd31^-/-^* mice (Figure 3B). Most metaphase spermatocytes underwent apoptosis and only few post-meiotic cells were detected in *Ankrd31^-/-^* mice (Figure S4J).

We conclude that compromised meiocyte survival provides a plausible explanation for fertility defects in *Ankrd31^-/-^* mice.

### Compromised synapsis in *Ankrd31^-/-^* meiocytes

Persistent DSBs and defective synapsis induce oocytes elimination in newborn mice and spermatocyte elimination in pachytene (Barchi et al., 2005; Burgoyne et al., 2009; Di Giacomo et al., 2005; Mahadevaiah et al., 2008; Royo et al., 2010; Wojtasz et al., 2012). In contrast, deficiencies in maturation of DSBs into COs lead to chromosome alignment defects and an arrest in metaphase I in spermatocytes (Burgoyne et al., 2009; Vernet et al., 2011). Hence, the complex patterns of elimination of *Ankrd31^-/-^* meiocytes prompted us to investigate meiotic recombination in *Ankrd31^-/-^* mice. Both chromosome axis formation and SC formation are used to stage meiotic prophase and provide a reference for kinetic studies of recombination (Table S1). Hence, we tested firstly if chromosome axis, and secondly if SC form with normal kinetics in *Ankrd31^-/-^* spermatocytes. Axes formed with wild type kinetics (Figure S5A), but SC formation was delayed on autosomes in *Ankrd31^-/-^* spermatocytes (Figure 3D and 3E). Despite this delay most *Ankrd31^-/-^* spermatocytes seemed to complete autosomal SC formation in adults (Figure 3D). Due to methodological limitations it is unclear if all spermatocytes succeeded in autosomal synapsis in adults (see STAR Methods for explanation). Autosomal SC formation occurred with high fidelity as we did not observe obvious non-homologous interactions in pachytene stage spermatocytes (n>200) in our *Ankrd31^-/-^* mutant line. SC formation was compromised in *Ankrd31^-/-^* oocytes (Figure S5B), which also manifested in higher rates of asynapsis in *Ankrd31^-/-^* oocytes (54.76%, n=126 oocyte) than wild type oocytes (22.96%, n=135 oocyte) at a fetal developmental stage where oocytes are in mid pachytene (Table S5). Together, these observations suggest that whereas ANKRD31 is not required for homology search *per se,* ANKRD31 is required for efficient and timely engagement of homologous chromosomes genome wide in both sexes.

Compared to autosomes, we observed a more penetrant defect in SC formation between the PARs of the heterologous sex chromosomes in males. PARs failed to synapse in 99.7% (n>420) and 99% (n=191) of pachytene spermatocytes from adult or juvenile *Ankrd31^-/-^* mice, respectively. Beyond asynaptic PARs, various abnormal sex chromosome configurations were observed in *Ankrd31^-/-^* spermatocytes (Figure 3F). This suggests that ANKRD31 plays a more important role in ensuring recombination between the PARs of heterologous sex chromosomes than autosomes.

### Delayed formation and resolution of early recombination intermediates in *Ankrd31^-/-^* meiocytes

To examine the possible reasons that underpin deficiencies in synapsis formation in *Ankrd31^-/-^* meiocytes we monitored kinetics of recombination intermediates (Figure 4 and S5C). Singlestranded DNA ends resulting from DSBs are marked by recombinases RAD51 and DMC1 and the RPA complex (Hunter, 2015; Moens et al., 2007). Levels of these markers on DSB repair intermediates change as recombination progresses. Whereas RAD51 and DMC1 tend to mark intermediates in unsynapsed regions (prevalent in leptotene and early zygotene), RPA accumulates to higher levels on intermediates in synapsed regions (prevalent in late zygotene and early pachytene). Focus numbers of all three markers drop as wild type meiocytes progress to late pachytene, indicating completion of DSB repair at this stage. RAD51, DMC1 and RPA foci accumulated with a delay in *Ankrd31^-/-^* spermatocytes (Figure 4A-E and S5C). Focus numbers remained abnormally low until early zygotene, increasing only after chromosome axes fully formed (late zygotene-like stage). Thereafter, RAD51, DMC1 and RPA foci persisted in abnormally high numbers until late prophase in *Ankrd31^-/-^* spermatocytes indicative of a repair defect (Figure 4C-E). To assess kinetics of DSB formation and repair we also measured another DSB marker, serine 139-phosphorylated histone H2AX (γH2AX) (Figure 4F and 4G). γH2AX accumulation is thought to mainly reflect DSB-dependent ATM activity in early prophase, and DSB- and unsynapsed axes-dependent ATR activity after mid zygotene (Barchi et al., 2008; Bellani et al., 2005; Royo et al., 2013). γH2AX levels were much lower in *Ankrd31^-/-^* than wild type spermatocytes in leptotene and early zygotene, indicative of lower ATM activity. In contrast, γH2AX robustly accumulated on unsynapsed chromatin from late zygotene, consistent with proficient ATR activation on unsynapsed chromatin. This also manifested in the apparently normal accumulation of γH2AX in the unsynapsed chromatin of sex chromosomes, called the sex body, in pachytene (Figure S5D). We also noted that higher numbers of γH2AX flares persisted on synapsed chromatin in late pachytene and diplotene spermatocytes in *Ankrd31^-/-^* mice as compared to wild-type (Figure S5D and S5E). These γH2AX flares are distinct from the sex body and likely indicate persisting recombination intermediates along synapsed sections of autosomes (Chicheportiche et al., 2007).

**Figure 4.**
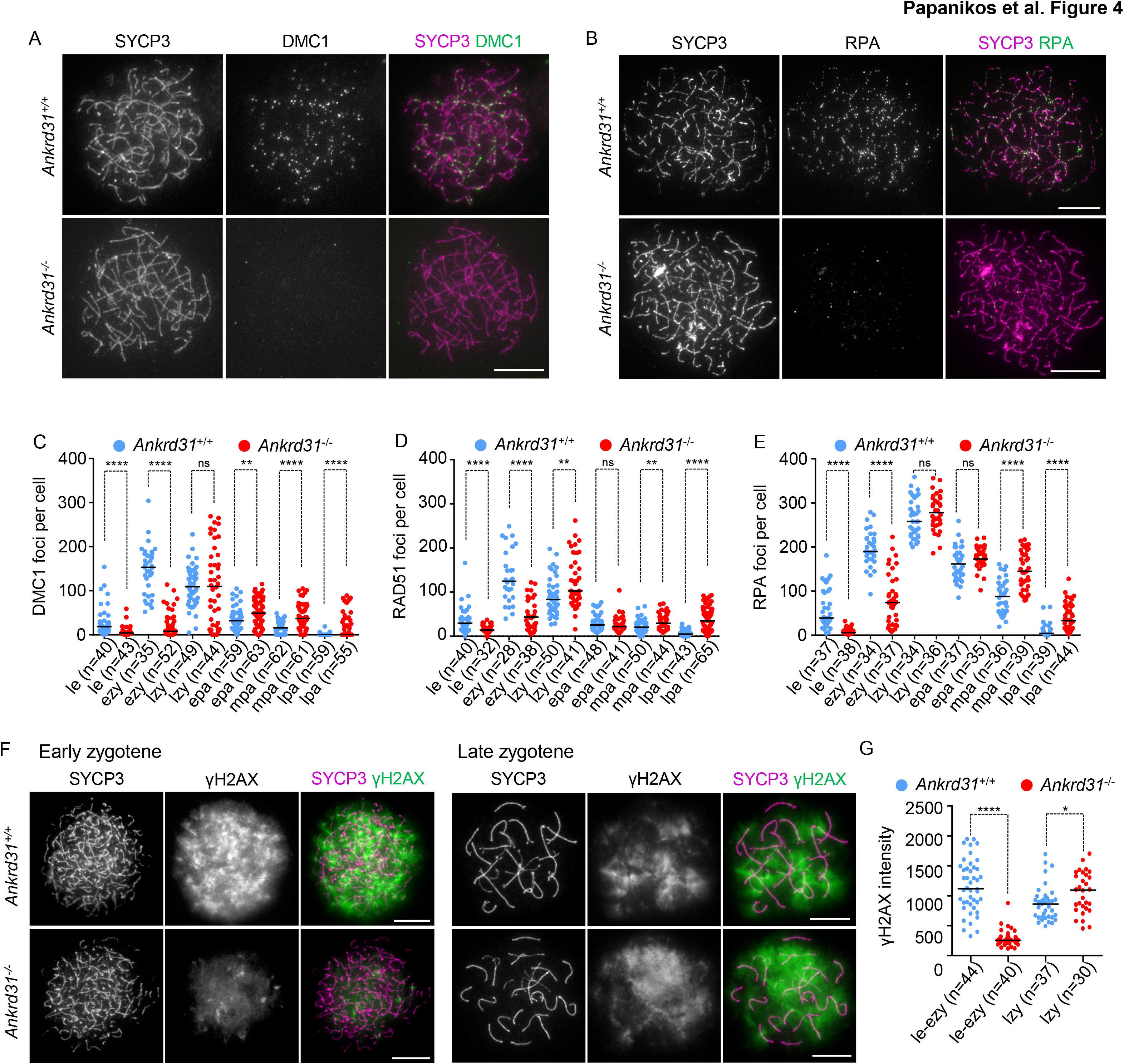
Delayed kinetics of recombination foci in *Ankrd31^-/-^* spermatocytes. (**A, B, F**) Indicated proteins were detected in nuclear surface spreads of spermatocytes. Bars, 10μm. Early zygotene cells are shown in **A** and **B**. Quantifications of DMC1 (**C**), RAD51 (**D**), and RPA (**E**) focus numbers and total nuclear γH2AX signal (**G**) in the indicated genotypes. (**C, D, E**) DMC1, RAD51 and RPA foci were quantified in leptotene (le), early zygotene (ezy), late zygotene (lzy), early pachytene (epa), mid pachytene (mpa) and late pachytene (lpa). (**G**) γ2AX signal was measured in pools of leptotene and early zygotene (le-ezy) or late zygotene (lzy) spermatocytes, levels are shown in each cell in arbitrary units. (**C, D, E, G**) Numbers (n) of counted cells and medians are indicated. Mann-Whitney U test calculated significance, nonsignificant (ns), 0.05>P>0.01 (*), 0.01>P>0.001 (**) and P0.0001 (****). See also Figure S5.

ANKRD31 had a qualitatively similar effect on the dynamics of DSB repair proteins in oocytes as compared to spermatocytes (Figure S5F-O). Accumulation of RAD51, DMC1 and RPA foci was delayed, and both RPA and γH2AX signal persisted longer in late prophase in *Ankrd31^-/-^* oocytes as compared to wild-type.

Overall, our data suggest that accumulation of DSB recombination intermediates is delayed in both sexes of *Ankrd31^-/-^* meiocytes, which delays and/or compromises SC formation. Persistence of recombination foci also indicates defective and/or delayed DSB repair. These defects provide a plausible explanation for elevated apoptosis in pachytene spermatocytes and oocytes of newborn mice. The DSB repair and SC defects appear subtle enough to allow the production of a cohort of oocytes for reproduction, which however cannot maintain high fertility with increasing age. Likewise, the subtlety of the DSB repair defects could explain the survival of a significant cohort of *Ankrd31^-/-^* spermatocytes beyond prophase.

### COs are lost from PAR in *Ankrd31^-/-^* spermatocytes

A penetrant defect in the PAR synapsis in spermatocytes (see Figure 3F) prompted us to test if a CO formation defect in X and Y PAR could account for the lagging chromosomes in metaphase arrested *Ankrd31^-/-^* spermatocytes (Figure 3B). Hence, we examined if *Ankrd31^-/-^* spermatocytes are defective in the formation of MLH1 marked recombination foci, which are thought to be precursors of ~90% of COs in mid and late pachytene (Figure 5A and 5B) (Gray and Cohen, 2016). Whereas MLH1 foci formed on autosomes in normal numbers in most *Ankrd31^-/-^* spermatocytes, MLH1 foci were always absent from the PARs of sex chromosomes in pachytene cells. We also examined chromosome spreads of metaphase spermatocytes. While in wild-type chiasmata were present between each pair of homologs and between sex chromosomes (n=79 cells), no *Ankrd31^-/-^* spermatocytes had chiasma between X and Y chromosomes as detected by FISH (n=57 cells, Figure 5C). A small fraction of cells (10.12% of 89 cells examined without FISH) also lacked chiasma between an autosome pair. Overall, these observations suggest that MLH1 foci form and mature with high efficiency on autosomes, but never on PARs, in spermatocytes. The loss of PAR-associated COs explains the observed metaphase arrest phenotype of *Ankrd31^-/-^* spermatocytes. Hence, it is likely that loss of ANKRD31 affects fertility more in males than females because ANKRD31 is required for recombination between male-specific heterologous sex chromosomes.

**Figure 5.**
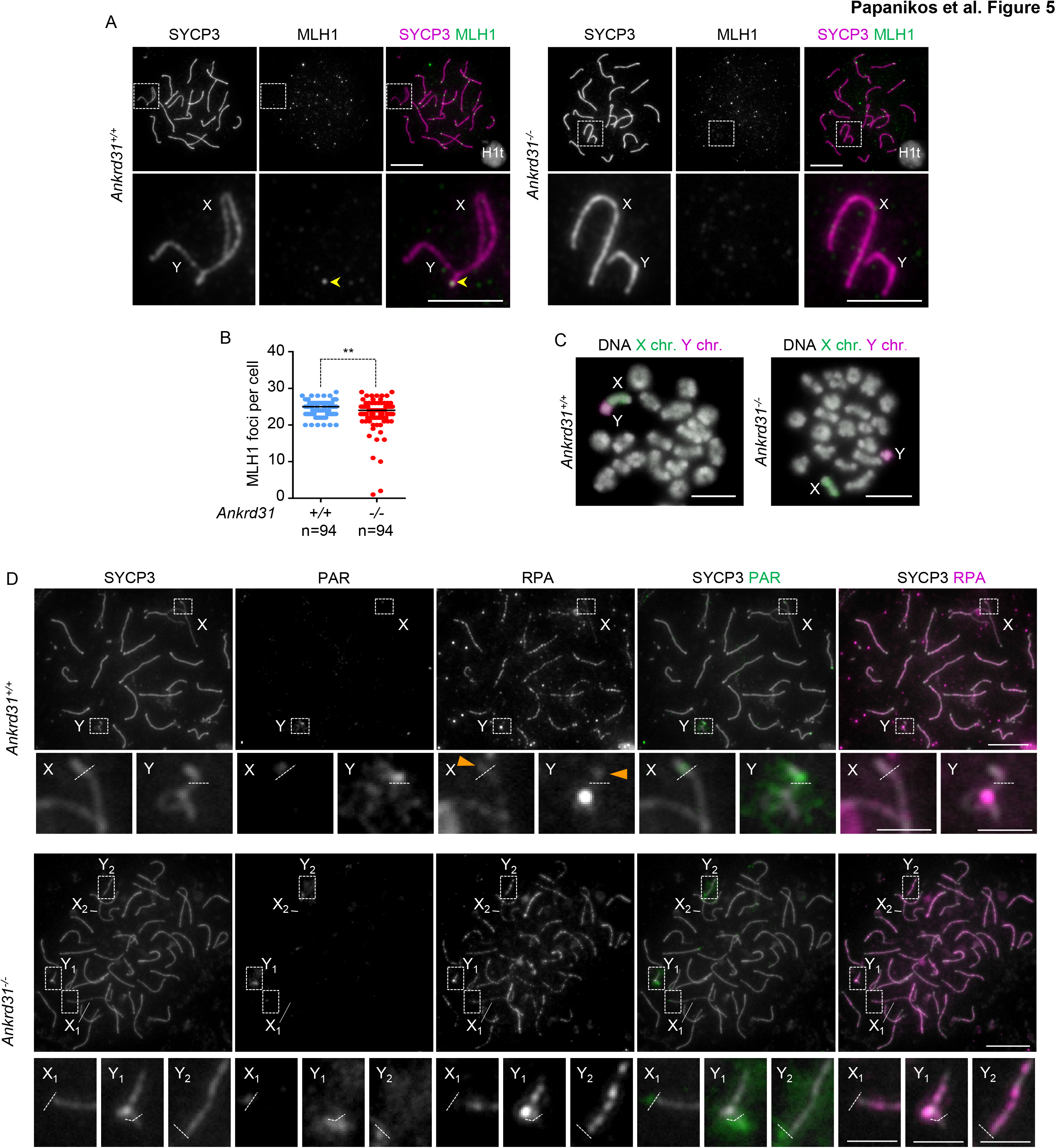
PARs fail to engage in *Ankrd31^-/-^* spermatocytes. (**A**) Indicated proteins were detected in nuclear spread *Ankrd31^+/+^* and *^-/-^* spermatocytes. Mid pachytene cells were identified by histone H1t staining (see miniaturized images of H1t stain in bottom right corner of overlay images). X and Y chromosomes are shown in enlarged insets in the bottom panel. PAR-associated MLH1 focus (yellow arrowhead) is marked in wild-type. (**B**) Quantifications of MLH1 focus numbers in the indicated genotypes. Numbers (n) of counted cells and medians are indicated. Significance was calculated by Mann-Whitney U test at 0.01>P>0.001 (**). (**C**) FISH detected X and Y chromosomes in metaphase spreads in indicated genotypes, DAPI detected DNA. (**D**) PAR FISH was combined with immunostaining of SYCP3 and RPA in surface spread spermatocytes. One late zygotene *Ankrd31^+/+^* and two early pachytene *Ankrd3^-/-^* are shown. Enlarged images of boxed areas are shown below their respective full nuclei images. Chromosome X and Y are indicated, white dashed line marks the boundary between the PAR FISH signal and the rest of the chromosome. Orange arrowhead points at axis-associated RPA foci within or distal to the PAR FISH signal. Enlarged images of Y chromosome PARs are shown for both *Ankrd31^-/-^* cells (Y_1_ and Y_2_). Enlarged image of X chromosome PAR is only shown from one of the cells (X_1_), because the X PAR overlays an autosomal axis in the second cell (X_2_). Bars are 10μm in low resolution images of (**A, C, D**), and 5μm or 2.5μm in enlarged insets in **A** and **D**, respectively.

CO formation may fail in PAR regions due to an absence of DSBs or a repair defect of existing DSBs. To test these possibilities we detected DSB repair foci (RPA) in combination with a FISH probe that recognizes PAR sequences that border the heterologous parts of X and Y chromosomes (Figure 5D). RPA foci were present on at least one of the two PARs in most wild type spermatocytes (91.7%, n=169) but only in a minority of (20.9%, n=67) *Ankrd31^-/-^* spermatocytes in late zygotene or early pachytene. This observation contrasts globally normal RPA focus numbers in *Ankrd31^-/-^* spermatocytes in late zygotene and early pachytene (Figure 4E). Thus, ANKRD31 seems to be more critical for DSB formation on PARs than in the rest of the genome.

### ANKRD31 is required for the normal spatial distribution of meiotic DSBs

The disproportionately severe reduction of DSB repair focus numbers in PARs as compared to autosomes suggested that the spatial distribution of DSB formation is altered in the absence of ANKRD31. To further test this possibility we mapped the genome-wide distribution of singlestranded DNA ends that result from DSB formation in testes of adult mice. We carried out chromatin-immunoprecipitation of the meiosis-specific recombinase, DMC1, and analyzed the immunoprecipitates by sequencing (ssDNA ChIP-seq, SSDS) (Khil et al., 2012; Smagulova et al., 2011) (Figure 6, S6 and Table S6, S7). SSDS informs on the relative steady state levels and genomic distribution of DMC1-associated single-stranded DNA in early recombination intermediates, which are a function of both DSB formation and the half-life of early recombination intermediates.

**Figure 6.**
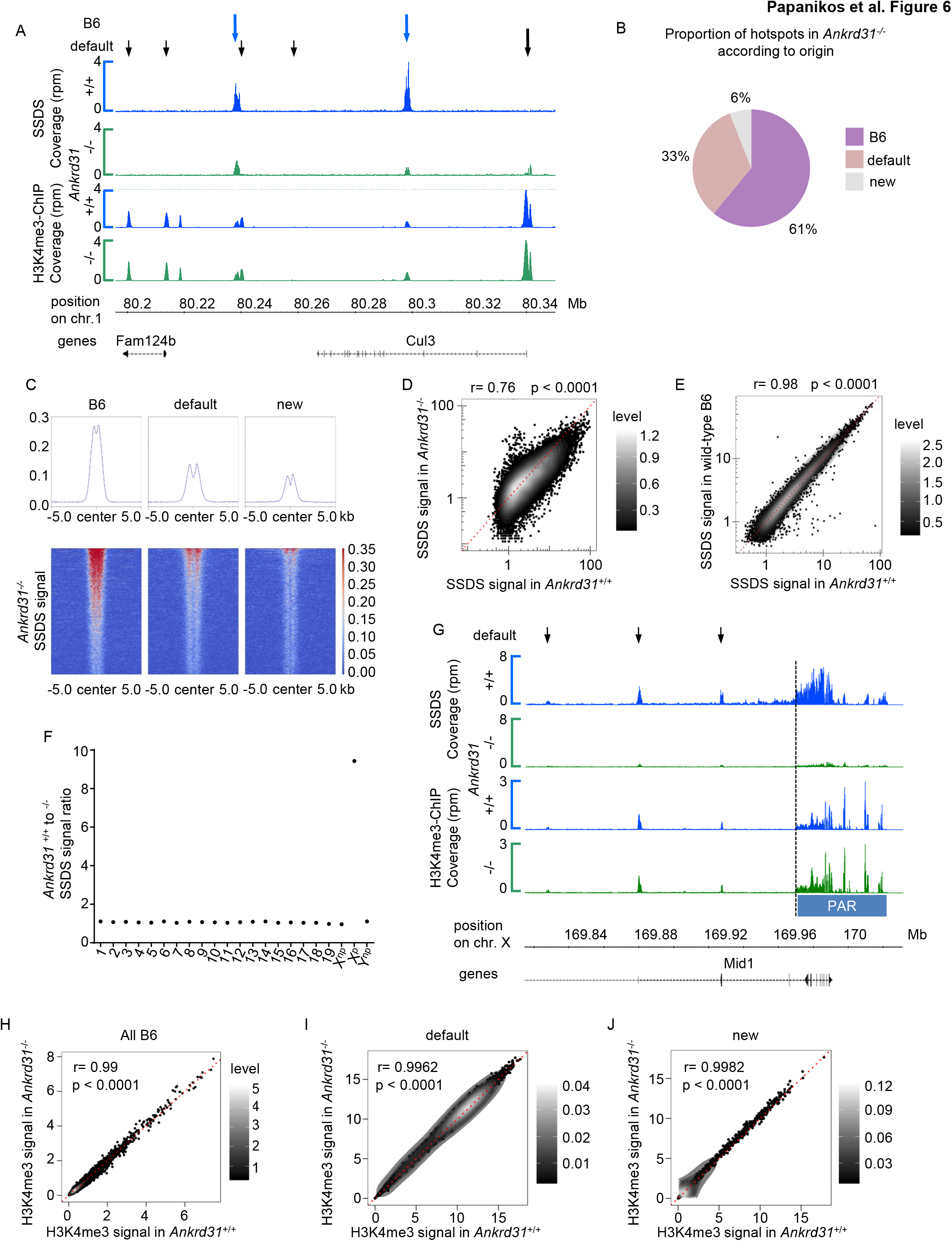
Spatial distribution of DSBs is altered in the absence of ANKRD31. (**A, G**) DMC1 ChIP SSDS and histone H3K4me3 signal coverage are shown in the indicated mouse lines in a 150kb region of chromosome 1 in **A** and in the broader PAR region of X chromosome in **G**. Genomic positions and annotated protein-coding genes are indicated. Coverage was merged from two experiments and normalized to library size. (**A**) Blue and black arrows mark sites of published C57BL/6J (B6) hotspots and PRDM9-independent default hotspots from Diagouraga et al. 2018 and Brick et al. 2012, respectively. Long and short black arrows represent comparatively strong or weak hotspots in *Prdm9^-/-^* mice according to DMC1 SSDS coverage. (**B**) Proportion of *Ankrd31* hotspots that match PRDM9-defined (i.e. B6) or default hotspots, or that do not match either (new). (**C**) Average plots (top) and heatmaps (bottom) of DMC1 SSDS signal in *Ankrd31^-/-^* mice at hotspots overlapping with B6, with default (i.e. from *Prdm9^-/-^*) or with neither of those (new). DMC1 read coverage was computed around the DMC1 peak center (±5.0kb). (**D, E**) SSDS signal comparison at PRDM9-defined hotspots between *Ankrd31^+/+^* and *Ankrd31^-/-^* in **D**, or between *Ankrd31^+/+^* and wild-type C57BL/6J mice (data from Diagouraga et al. 2018) in **E**. (**F**) *Ankrd31^+/+^* to *^-/-^* DMC1 SSDS signal ratios are shown for each autosome, the non-PAR regions of X (X^np^) and Y (Y^np^), and available sequence of the X PAR. (**G**) Arrows mark PRDM9-independent hotspots in the non-PAR region upstream of PAR. (**H-J**) H3K4me3 signal comparison at PRDM9-defined (B6) (**H**), default (**I**) or new (**J**) hotspots sites between *Ankrd31^+/+^* and *Ankrd31^-/-^* mice. H3K4me3 read coverages were normalized as described in the STAR Methods. (**D, E, H-J**) Dotted red lines represent one to one relationship between x and y. Levels represent the density calculated with the density2d function from R. Pearson s correlation coefficient (r) is indicated with significance (p). See also Figure S6, Table S6 and S7.

### An increased usage of default hotspots

Hotspot numbers were higher in *Ankrd31^-/-^* (22215) than wild type control (13507) testes. We also observed major differences in the spatial distribution of DSB hotspots between *Ankrd31^-/-^* testis and wild-type (Figure 6A). Whereas the majority of *Ankrd31^-/-^* DSB hotspots (61%) overlapped with PRDM9-dependent hotspots of wild-type (hereafter called B6 hotspots), a significant fraction (33%) overlapped with previously described default hotspots of *Prdm9^-/-^* mice that mostly associate with active promoters (Figure 6A, 6B) (Brick et al., 2012). A further 6% of the *Ankrd31^-/-^* hotspots were at previously undescribed sites (hereafter called “new”). In *Ankrd31^-/-^*, the average strength of hotspots overlapping with B6 hotspots was the highest and “new” hotspots were the weakest (Figure 6C). Hence, default and “new” hotspots contributed less to the total single-stranded DNA signal than what would be expected based on their numbers (compare Figure 6B and S6A).

In addition, a redistribution of the intensity of SSDS signals was observed at hotspots overlapping with B6 hotspots in *Ankrd31^-/-^* testes. This was illustrated by a lower correlation in hotspot-associated SSDS signals (r=0.76, Pearson) between *Ankrd31^-/-^* and wild-type than between two wild type samples (r= 0.98) (Figure 6D and 6E). Consistent with an apparent redistribution of single–stranded DNA signal, not all the B6 hotspots were detected as hotspots in *Ankrd31^-/-^* testes. Weak B6 hotspots were detected in lower proportions in *Ankrd31^-/-^* testes than strong B6 hotspots (Figure S6B). Likewise, strong default hotspots of *Prdm9^-/-^* mice emerged as hotspots in *Ankrd31^-/-^* mice with higher frequency than weak default hotspots (Figure S6C). The combination of these observations suggested that there was a general shift from the use of PRDM9-dependent B6 hotspots towards default hotspots in *Ankrd31^-/-^* mice. We conclude that there is a differential hotspot usage and/or DSB repair activity in *Ankrd31^-/-^* compared to wild type which could be also influenced by altered timing of recombination initiation and/or SC formation.

### A strong reduction of DSB activity in the PAR

Given that loss of ANKRD31 altered the distribution of SSDS signal at the hotspot level, we wondered if ANKRD31 may also have differential effects in specific chromosomes and in the PAR. Hence, we calculated wild-type to *Ankrd31^-/-^* SSDS signal ratio for each chromosome and for the sequenced portion of the PAR (Figure 6F, see also STAR Methods for mapping reads to PAR). Ratios of SSDS signals were within +/- 9 % of genome average on each autosome and sex chromosomes regions excluding the PAR. In contrast, SSDS coverage was strongly reduced (9.4 fold) at the PAR in *Ankrd31^-/-^* testes relative to wild-type (Figure 6F, 6G and Table S7). SSDS coverage was also strongly reduced in hotspots within a 150 kb region upstream of X PAR (Figure 6G and Table S7).

These observations support the hypothesis that ANKRD31 has a particularly critical role in recombination and DSB formation at PARs.

### Histone H3K4me3 distribution on chromatin is similar in wild type and *Ankrd31^-/-^* testes

Global changes in the timing and the spatial distribution of DSBs in *Ankrd31^-/-^* mice could be caused by a change in the testicular transcriptome or altered chromatin organisation. However, we detected very few, if any, changes in the testis transcriptome of juvenile (12 days old) *Ankrd31^-/-^* mice relative to wild-type (Figure S6D and S6E). Testicular PRDM9 protein levels were also similar in *Ankrd31^-/-^* and wild type testes (Figure S6F). Both PRDM9-dependent and PRDM9-independent hotspots are characterized by enrichment for histone H3K4me3, therefore we performed histone H3K4me3 ChIP-seq on testes of juvenile mice (12 days old) to test if an altered distribution of H3K4me3 could explain the patterns of DSB formation in *Ankrd31^-/-^* mice (Figure 6A and 6G). To allow better comparison of different samples we normalized H3K4me3 signal to a common set of transcription start sites (Davies et al., 2016 and STAR Methods).

H3K4me3 signal enrichment appeared highly similar in wild type and *Ankrd31^-/-^* mice both at B6 hotspots and PRDM9-independent hotspots including the broader region of the PAR (Figure 6A, 6G, and Table S7). Indeed, histone H3K4me3 enrichment strongly correlated between wild type and *Ankrd31^-/-^* testes at B6 (r=0.99) and default (r=0.99) hotspot sites (Figure 6H, 6I) indicating that PRDM9-dependent and -independent H3K4me3 deposition is not affected in *Ankrd31^-/-^* mice. Furthermore, we found no significant correlation between the variation of wild-type to *Ankrd31^-/-^* SSDS signal ratios and H3K4me3 signal ratios at B6 hotspots (Figure S6G).

Together, these observations suggest that ANKRD31 is required for neither PRDM9 methyltransferase activity at B6 hotspots nor the accumulation of H3K4me3 at PRDM9- independent hotspots including the PAR. The shift in DSB activity from B6 to default hotspots and the severe reduction of DSB activity in the PAR in *Ankrd31^-/-^* mice are therefore not due to changes in histone H3K4me3 levels.

### Timely assembly of DSB-promoting proteins into chromatin-bound complexes depends on ANKRD31

Given its localization on axes, ANKRD31 might modulate the function of pre-DSB recombinosomes, which could provide a feasible explanation for *Ankrd31^-/-^* phenotypes. Consistent with this hypothesis we found that both MEI4 and REC114 foci appeared with delayed kinetics in *Ankrd31^-/-^* spermatocytes (Figure 7A-C and S7A-C). Nevertheless, MEI4 and REC114 foci did accumulate by zygotene and persisted until SC formation was complete. Kinetics of MEI4 and REC114 focus formation matched those of early recombination intermediates, suggesting that a delay in the formation of MEI4- and REC114-marked pre-DSB recombinosomes contributes to a delay in DSB formation in *Ankrd31^-/-^* meiocytes.

**Figure 7.**
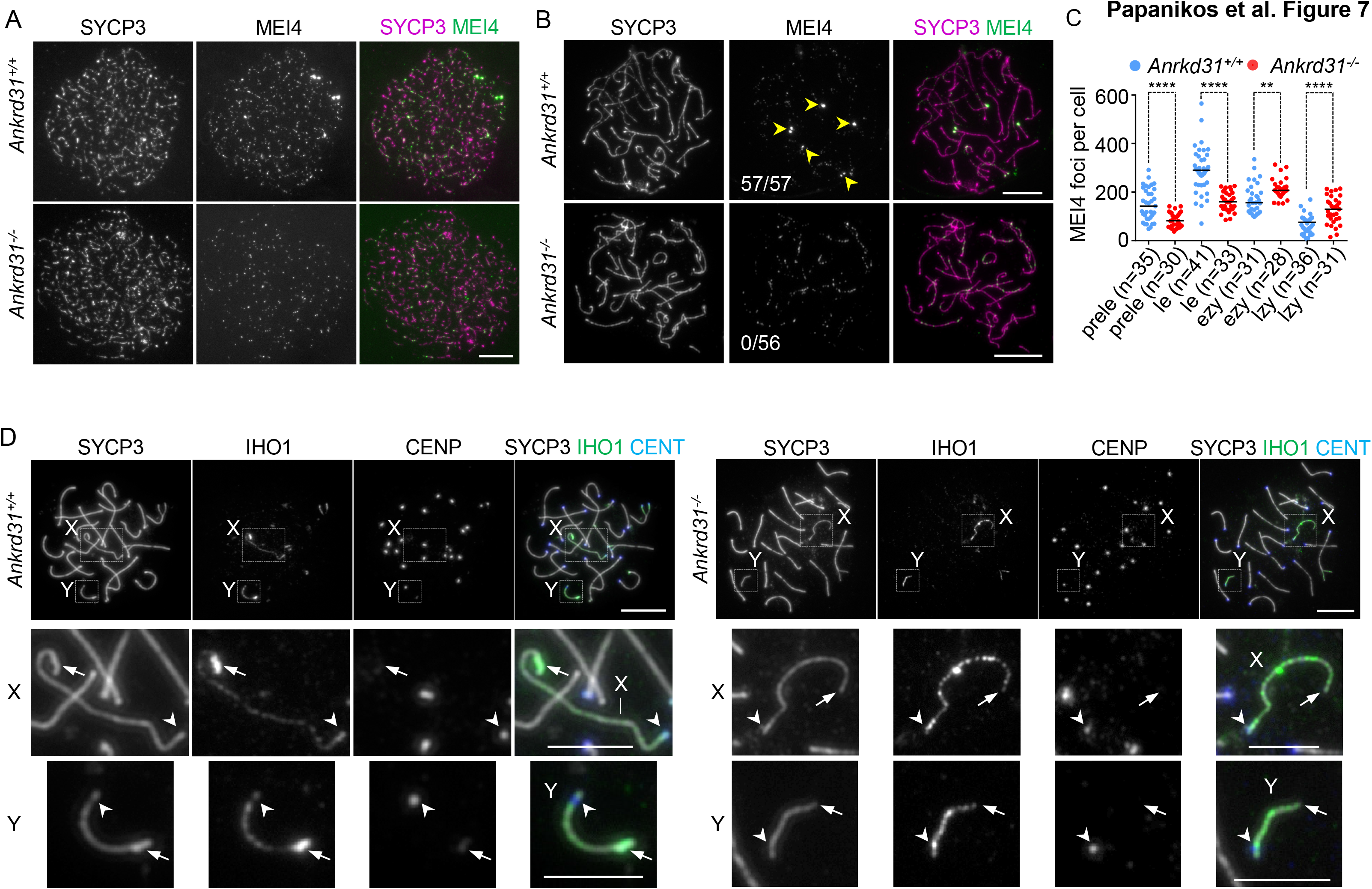
ANKRD31 is required for timely formation of MEI4 foci and the formation of modified PAR axis that is rich in DSB-promoting proteins **(A, B, D)** Immunofluorescence detected indicated proteins and centromere (CENT in **D**) in nuclear spread spermatocytes of indicated genotypes. Leptotene (**A**) and late zygotene (**B, D**) spermatocytes are shown. (**A, B**) Exposure of MEI4 was matched between *Ankrd31^+/+^* and ^-/-^genotypes. Yellow arrowheads mark MEI4 aggregates at the ends of 5 chromosomes. The fraction of late zygotene spermatocytes that contained MEI4 aggregates is indicated in the MEI4 image. (**C**) Quantification of MEI4 focus numbers in preleptotene (prele), leptotene (le), early (ezy) and late (lzy) zygotene spermatocytes of indicated genotypes. Numbers (n) of examined cells and median focus counts are indicated. Mann-Whitney U test calculated significance at 0.01>P>0.001 (**) and P<0.0001 (****). (**D**) Enlarged images of boxed areas around X and Y chromosomes are shown in the bottom two panels for each genotype. PAR (arrow) and centromeric (arrowhead) ends are marked. Bars are 10μm and 5μm in low and high magnification images, respectively, in **A**, **B** and **D.** See also Figure S7.

### Robust aggregates of DSB-promoting proteins in PARs depend on ANKRD31

Whereas MEI4 and REC114 foci eventually formed in *Ankrd31^-/-^* meiocytes, we did not observe MEI4 and REC114 aggregates in PARs or autosome ends (Figure 7B and S7B). We also noticed an alteration of chromosome axis at PARs in *Ankrd31^-/-^* meiocytes. SYCP3-staining appeared brighter and/or thicker at the PAR-end of sex chromosomes (particularly on Y chromosome) in late zygotene and at the zygotene-to-pachytene transition in wild-type (Figure 1E, 7D and S7D). The unsynapsed axis-binding IHO1 protein, which is thought to form the main interface between axes and pre-DSB recombinosomes, was also always markedly enriched on the thickened PAR axes in wild-type spermatocytes (Figure 7D and S7D). This enrichment of IHO1 matched formation of MEI4 and REC114 aggregates (Figure 2G and S3C), and also persisted in the PARs after synapsis initiation (Figure S7D and S7E). In *Ankrd31^-/-^* spermatocytes, IHO1 robustly decorated unsynapsed axes, but its PAR-enrichment was strongly reduced; only a weak enrichment of IHO1 could be discerned in a small minority of late zygotene and zygotene-to-pachytene transition cells (15.56%, n=59 cells, n=2 mice). Mirroring this change, the bright/thickened SYCP3 stain was abolished at the PAR in *Ankrd31^-/-^* spermatocytes. Together, these observations suggest that ANKRD31 is required for the establishment of a specialized PAR axis domain that is particularly rich in DSB-promoting proteins.

## Discussion

The prevailing model of DSB formation posits that pre-DSB recombinosomes recruit open chromatin to chromosome axes to promote DSB formation (Grey et al., 2018; Panizza et al., 2011). We report that ANKRD31 co-regulates spatial and temporal patterns of pre-DSB recombinosome assembly and DSB activity, which supports the prevailing model and provides new insight into the regulatory pathways that underlie DSB control. Our findings are supported by similar observations in the accompanying manuscript by Boekhout et al. 2018.

### ANKRD31 is part of the pre-DSB recombinosome

Using transcriptome data, we identified ANKRD31 as a gene with possible roles in meiotic recombination. Consistent with this, we find that ANKRD31 colocalizes with highly conserved protein components of pre-DSB recombinosomes (Figure 2, S3) (IHO1, MEI4 and REC114 in mice) (Arora et al., 2004; de Massy, 2013; Kumar et al., 2010; Kumar et al., 2018; Stanzione et al., 2016). Although the precise architecture of these complexes is unknown, ANKRD31 association with pre-DSB recombinosomes seems to involve direct interactions with REC114 and IHO1 (Figure 2, S3 and Boekhout et al. accompanying manuscript). Surprisingly, we observed two types of pre-DSB recombinosomes in mouse meiocytes. One type is detected as foci along chromosome axes as observed in previous studies (Kumar et al., 2010; Stanzione et al., 2016), the second type is detected as large aggregates on the PAR and the ends of three autosomes. Acquaviva et al. discovered that the distal ends of chromosome 4, 9 and 13 share homology with PAR and serve as sites of autosomal pre-DSB recombinosome aggregates (personal communication from Acquaviva, Jasin & Keeney, which we reconfirmed using FISH, unpublished data). We refer collectively to these autosomal sites as PAR-like regions. Pre-DSB recombinosome foci and aggregates have distinct properties. Focus formation requires IHO1 and is enhanced by, but not dependent on, ANKRD31. In contrast, aggregate formation on PARs is independent of IHO1 but requires ANKRD31. IHO1 is thought to recruit other components of pre-DSB recombinosomes to axes in most parts of the genome through its interaction with HORMAD1. Hence, we propose that the mode of pre-DSB recombinosome recruitment to the axis is altered in the PAR and PAR-like autosomal regions. MEI4, REC114 and ANKRD31 are mutually dependent for aggregate formation at the PAR (Figure S3), and ANKRD31 might be a specificity-factor that recognizes some properties of these regions to allow recruitment of pre-DSB recombinosome components. Alternatively, ANKRD31 might confer specific properties on pre-DSB recombinosome components, enabling them to be enriched in the unique chromatin environment of PAR and PAR-like regions. We noted that the noticeable thickening of the axis in the PAR depends on ANKRD31 (Figure 7D). The formation of a disproportionately long PAR axis, short DNA loops and extensive axis splitting, which underlies the axis thickening in our study, all depend on ANKRD31 (Acquaviva, Jasin & Keeney, personal communication). Whether ANKRD31 reorganizes PAR chromatin directly or indirectly (via MEI4 or REC114) is unknown.

### ANKRD31 may control recombination initiation by regulating pre-DSB recombinosomes

The effect of ANKRD31 on pre-DSB recombinosome formation raises the question of how ANKRD31 controls recombination initiation. Using two complementary approaches to assess recombination initiation, our study revealed two clear consequences of the absence of ANKRD31. First, a delay in DSB repair focus formation and, second, a change in the localization and relative levels of DSBs along chromosomes, with the most noticeable consequences in the PAR. The delay in the accumulation of early recombination intermediates (DMC1, RAD51 and RPA foci) was consistent with the delayed appearance of pre-DSB recombinosomes in *Ankrd31^-/-^* spermatocytes. We thus propose that ANKRD31 enhances pre-DSB recombinosome assembly on chromosome axes, which helps timely initiation of recombination and SC formation between homologs (Figure S7E). A related, but possibly distinct, molecular function of ANKRD31 may serve to permit the aggregate formation of DSB-promoting proteins in PAR. It is tempting to speculate that ANKRD31 ensures obligate recombination initiation in the PAR by allowing the formation of these aggregates (Figure S7E).

### Altered DSB localization to PRDM9-independent sites in *Ankrd31^-/-^* mice

DSB activity at PRDM9-independent default sites is normally repressed in wild-type by a poorly understood PRDM9-dependent mechanism, which may reflect competition between PRDM9-dependent and independent sites (Brick et al., 2012; Diagouraga et al., 2018). This effect seems important, because extensive DSB activity at default sites is thought to cause asynapsis and delay in DSB repair (Brick et al., 2012; Hayashi et al., 2005). One of the striking features of the change in DSB localization in *Ankrd31^-/-^* mice is the use of default sites, which could contribute to the synapsis defect and delayed DSB repair kinetics in *Ankrd31^-/-^* mice. It is unlikely that use of default sites is due to changed PRDM9 activity in *Ankrd31^-/-^* meiocytes, since histone H3K4me3 distribution was not significantly changed (Figure 6, S6 and Table S7). We favor the idea that default site usage is a consequence of altered pre-DSB recombinosome regulation. ANKRD31-dependent properties of pre-DSB recombinosome could impact upon competition between PRDM9-dependent and independent sites. Alternatively delayed DSB formation might by itself promote the usage of default sites in *Ankrd31^-/-^* mice. A potential link between timing of DSB formation and usage of PRDM9-independent sites has been previously observed; PAR-associated DSBs are mostly independent of PRDM9 and form later than autosome-DSBs (Kauppi et al., 2011).

### ANKRD31 may impact ATM activity

Boekhout et al. (in accompanying manuscript) estimated total DSB activity in testes of *Ankrd31^-/-^* mice by measuring levels of covalently linked SPO11-oligonucleotide complexes (SPO11-oligo), which are generated during the normal processing of DNA ends of DSB sites (Neale et al., 2005). These measurements indicated two-to-four fold higher DSB activity in testes of *Ankrd31^-/-^* mice as compared to wild-type. Curiously, a corresponding increase in DSB repair focus numbers was not observed in pre-pachytene *Ankrd31^-/-^* meiocytes (Figure 4 and Boekhout et al.). The persistence of DSB repair foci in late prophase in *Ankrd31^-/-^* meiocytes makes increased turnover of recombination intermediates unlikely to account for the disparity between SPO11-oligo and DSB repair focus measurements. Rather, this could be explained if multiple DSBs can form abnormally at adjacent sites in *Ankrd31^-/-^* mice. In both mouse and yeast, ATM inhibits the occurrence of multiple DSBs at adjacent sites both on the same chromatid and sister chromatids (Garcia et al., 2015; Lange et al., 2016; Zhang et al., 2011). Disruption of ATM activity leads to elevation of DSB formation without a corresponding increase in repair foci (Barchi et al., 2008; Lange et al., 2011; Widger et al., 2018). Although we did not undertake a specific analysis of ATM activity, the strong reduction in γH2AX in early prophase spermatocytes is consistent with reduced ATM activity in *Ankrd31^-/-^* mice. Given the known functions of ATM in promoting DNA end resection and DSB repair (Barchi et al., 2008; Bellani et al., 2005; Joshi et al., 2015; Mimitou et al., 2017), a putative defect in ATM activation by DSBs could also contribute to delayed kinetics of DSB repair foci and synapsis in *Ankrd31^-/-^* meiocytes. Nonetheless, characteristic features of *Ankrd31^-/-^* meiocytes, i.e. loss of DSBs on PAR and elevated use of default hotspots (Figure 5 and 6), are not phenocopied by *Atm^-/-^* meiocytes (Lange et al., 2016). We therefore hypothesize that, in addition to regulating pre-DSB recombinosome formation, ANKRD31 permits robust ATM activation after DSB formation.

### DSB control on the PAR

The PAR region is highly enriched for DSBs compared to the genome average (Kauppi et al., 2011; Lange et al., 2016), and we found that this enrichment requires ANKRD31 (Figure 6 and Table S7). Intriguingly, DSBs were lost from the PAR in *Ankrd31^-/-^* mice independently of histone H3K4me3 enrichment (Figure 6 and Table S7). Enrichment of histone H3K4me3 had previously been linked to high DSB activity in the PAR (Brick et al., 2012) but is not sufficient to ensure high DSB numbers according to our results. We propose that robust DSB formation in the PAR requires an ANKRD31-dependent enrichment of pre-DSB recombinosomes (Figure 7 and S7) and/or ANKRD31-dependent alterations to axis/loop organisation as also suggested by (Kauppi et al., 2011) and personal communication from Acquaviva, Jasin & Keeney. Thus, the distribution of pre-DSB recombinosomes on axes seems to have a crucial role in defining genomic sites where DSBs occur.

Curiously, for unknown reasons, DSB activity is PRDM9-independent in the PAR in mice (Brick et al., 2012). We speculate that this might have arisen due to the erosion of PRDM9-binding sites during evolution. Such erosion occurs when the template used for the repair of DSBs at hotspots includes polymorphisms that disrupt PRDM9 binding sites (Lesecque et al., 2014; Myers et al., 2010). Since the strength of this effect increases with hotspot strength, high DSB activity in the PAR region is expected to erode PRDM9 sequence motifs faster than in other genomic regions. One way in which species could cope with the loss of PRDM9 binding sites is the concurrent evolution of the PRDM9 protein (Latrille et al., 2017). Another way is to use localized PRDM9-independent DSB initiation, which might require relieving PRDM9-dependent repression of default hotspot sites in this region, without affecting PRDM9 control in the rest of the genome. This could occur by delaying DSB formation in the PAR to a time-window where PRDM9 is inefficient in providing competitive advantage to PRDM9-dependent hotspot sites and/or blocking default hotspots. A non-exclusive mechanism could be the assembly of pre-DSB recombinosome aggregates with unique characteristics, which might permit the establishment of a PAR-specific chromatin environment where PRDM9-independent DSBs are possible. PRDM9- independent formation of DSBs in the PAR is not a universal feature, since there are PRDM9-dependent hotspots in human PARs (Pratto et al., 2014; Sarbajna et al., 2012). It will be interesting to determine whether the high level of accumulation of pre-DSB proteins is an evolutionary conserved feature of PARs, or if it is linked to PRDM9-independence in DSB formation. We predict that high accumulation of pre-DSB recombinosome proteins will promote high DSB density, regardless of the extent of PRDM9 involvement.

### Summary

The behavior of ANKRD31 and the phenotype of *Ankrd31^-/-^* meiocytes show that correct timing and spatial patterning of recombination initiation are dependent on ANKRD31 in mice. Our observations strongly suggest that ANKRD31 performs these functions by modulating the properties, formation kinetics and/or the stability of complexes of essential DSB-promoting proteins localized on chromosome axes. The phenotype of *Ankrd31^-/-^* mice provides support for the previously untested hypothesis that spatial patterning of recombination initiation is controlled not only by the chromatin environment of potential DSB sites in chromatin loops, but also by the DSB forming machinery that assembles in the topologically distinct compartment of the chromosome axis. This paradigm is best exemplified in the PARs of heterologous sex chromosomes, where obligate recombination seems to depend on ANKRD31-mediated enrichment of DSB promoting proteins and modification of chromatin organisation.

## ACKNOWLEDGEMENTS

We thank L. Acquaviva, M. Boekhout, M.E. Karasu, M. Jasin and S. Keeney for sharing critical unpublished results; D. Tränkner and M. Munzig for technical lab assistance and supporting mouse work, R. Jessberger (Institute of Physiological Chemistry, Faculty of Medicine at the TU Dresden, Germany) for sharing ideas, antibodies (anti-SYCP3) and departmental support; M. Stevense, G. Pearce and E. Hoffmann for discussion, and proofreading the manuscript; Transgenic Core Facility of Max Planck Institute of Molecular Cell Biology and Genetics (Dresden, Germany) for CRISPR injections and embryo transfers; Experimental Center of Medizinisch Theoretisches Zentrum (Dresden, Germany) for assisting in antibody production and taking care of mice; Deep Sequencing Facility of Center for Regenerative Therapies (TU Dresden, Germany) for RNA-sequencing and bioinformatics support; the MPI-CBG scientific computing facility for last minute bioinformatics support; I.R. Adams for collecting fetal ovaries and proofreading; I.R. Adams, I. MacGregor, A.M Pendes, L.G. Hernandez for testing usability of ANKRD31 antibodies in various mammalian species; K. Daniel (Biotechnology Center TU Dresden-BIOTEC, Germany) for ovaries and testis RNA preparation; Christine Brun for preparing material for immune-cytochemistry; Montpellier Resources Imagerie (MRI) for microscopy, the ENCODE Consortium and the ENCODE production laboratory(s) for the generation of mouse tissue transcriptome datasets. The Deutsche Forschungsgemeinschaft (DFG; grants: TO421/3-1/2, TO421/6-1/2, TO421/7-1, TO421/10-1, TO421/5-1, TO421/8-1/2) and HFSP research grant RGP0008/2015 supported F.P., I.D., R.R., S.V. and A.T. We thank the DIGS-BB Program for supporting A.B. J.C., C.G. and B.d.M. were funded by grants from the Centre National pour la Recherche Scientifique (CNRS) and the European Research Council (ERC) Executive Agency under the European Community’s Seventh Framework Programme (FP7/2007-2013 Grant Agreement no. [322788]). B.d.M. was recipient of the Prize Coups d’Élan for French Research from the Fondation Bettencourt-Schueller. E.T. and M.B. were funded by Telethon Foundation, (Grant GGP12189) and Mission Sustainability Grant, University of Rome Tor Vergata (Grant 141). D.L., P.J. and J.F. were funded by Czech Science Fundation (CSF): grant 13-08078S to J.F. Ministry of Education, Youth and Sports (MEYS): LQ project of the NSPII to J.F. A.S. was supported by Boehringer Ingelheim GmbH and Austrian Academy of Sciences.

## AUTHOR CONTRIBUTIONS

F.P. performed characterization of *Ankrd31M^-/-^* male phenotype, generated and validated ANKRD31 antibodies, generated *Ankrd31M^-/-^* mice, provided samples for ChIP-seq and RNA-seq experiments

J.C. performed ChIP-seq bioinformatics analysis

E. T. performed and analyzed PAR-FISH experiments

I. D. assisted in the characterization of *Ankrd31M^-/-^* male phenotype, performed ANKRD31 colocalization experiments, analyzed PAR-FISH experiments and ANKRD31 localization in *Mei4^-/-^* mice, validated ANKRD31 antibodies and produced H1t antibody R.R. performed *Ankrd31M^-/-^* female analysis

C.G. performed ChIPs, constructed libraries, purified PRDM9 antibody and measured PRDM9 protein levels in *Ankrd31^-l-^*

A.B. performed yeast two-hybrid experiments and immunoprecipitation of ANKRD31 in *Ankrd31^-/-^*

S.V.C. validated ANKRD31 antibodies, characterized ANKRD31 localization and assessed apoptosis in testis cryosections

A. S. performed multiple alignments of ANKRD31 protein sequences

M.S. performed initial Y2H experiments and initial IHO1 localization analysis at PAR P.J., D.L. and J.F. performed chromosome FISH experiments

F. J. produced CRISPR/CAS9 mRNA and assisted in gRNA preparation

M.B. analyzed PAR-FISH experiments, supervised E.T. and revised the manuscript

B. d.M. performed ANKRD31 localization experiments in *Mei4^-/-^* and *Rec114^-/-^* mice, analyzed data and co-wrote the section on ChIP-seq, introduction and discussion with A.T. and revised the rest of the manuscript.

A.T. designed experiments, analyzed data, supervised the project and wrote the manuscript with input from the other authors.

## Contact for Reagent and Resource Sharing

Further information and request for resources and reagents should be directed to the Lead Contact, Attila Toth.

## Methods

### Generation and genotyping of Ankrd31-knockout mice

*Ankrd31* mutant lines were generated using CRIPSR/Cas9 genome editing (Hwang et al., 2013; Shen et al., 2013; Wang et al., 2013), targeting exon6 of *Ankrd31* gene. A mixture of gRNA: AGT CACCAAAACACT GG (12.5 ng/μl) (designed using the online platform at http://crispr.mit.edu/) and Cas9 nuclease mRNA (50 ng/μl) was injected into pronucleus/cytoplasm of fertilized oocytes. The oocytes were subsequently transferred into pseudopregnant recipients. Injections and embryo transfer were performed by Transgenic Core Facility of MPI-CBG (Dresden, Germany). Out of 115 injected embryos, 82 were transferred into females and 15 were born. 14 of the 15 born pups had alterations in the targeted genomic locus. Two mice that were heterozygote for predicted frame-shift causing alleles (mut1 and mut2, Figure 3A) were bred with C57BL/6JCrl wild type mice to establish mouse lines. All experiments reported in the manuscript are based on samples from mice that were derivative of founder lines after at least three backcrosses.

#### gRNA production

The guide RNA (gRNA) expression vector DR274 (Addgene #42250) was used to construct the gRNA (Hwang et al., 2013). Primers encoding gRNA sequence were annealed to form doublestrand DNA with overhangs for ligation into DR274 vector that was linearized with Bsal restriction enzyme (NEB). PCR product was amplified from the resulting plasmid with DR274_F/DR274_R primers and used as a template for *in vitro* transcription with MEGAscript™ T7 Transcription Kit (Ambion).

#### CAS9 mRNA preparation

To prepare Cas9 mRNA, we first used the restriction enzyme PmeI to linearize the plasmid MLM3613 (Addgene #42251) (Hwang et al., 2013) that harbors a codon optimized Cas9 coding sequence and a T7 promoter for Cas9 mRNA *in vitro* synthesis. We then used the linearized MLM3613 as template to synthesize the 5’ capped and 3’ polyA-tailed Cas9 mRNA using the mMESSAGE mMACHINE^®^ T7 Ultra Kit (ThermoFisher, cat no: AM1345) according to the manufacturer’s instructions.

#### Genotyping protocol

Tail biopsies were used to generate genomic DNA by overnight protease K digestion at 55°C in lysis buffer (200mM NaCl, 100mM Tris-HCl pH 8, 5mM EDTA, 0.1% SDS). Following heat inactivation for 10 min at 95°C. These genomic preparations were used for PCR. F0 mice were genotyped by PCR amplification followed by PAGE electrophoresis and DNA sequencing. Mice in subsequent crosses were genotyped by PCR amplification followed by agarose gel electrophoresis. Combination of four primers, AS-79WT-FW2, Ank_ex5gen_LngR, Ank_ex5gen_LngFW, KO79-RV, were used to genotype the *Ankrd31^mut1^* allele. PCR product sizes were 623bp and 262bp for wild type, and 624bp and 400bp for *Ankrd31^mut1^* allele. Two primers, Ank_ex5_gen_shFW and Ank_ex5_gen_shR, were used to genotype *Ankrd31^mut2^* allele. PCR product sizes were 281bp for wild type, and 248bp for *Ankrd31^mut2^* allele.

### Animal experiments, choice of adults or juveniles

Gonads were collected from mice after euthanasia. Most cytological experiments of spermatocytes were carried out on samples collected from adult mice unless indicated otherwise. In particular, we used juvenile mice (13-14 days old) to enrich for cells that are in late zygotene or zygotene-to-pachytene transition. Juvenile mice were also used for transcriptome analysis and histone H3K4me3 ChIPseq, where differences between the cellularities of testes in adult wild type and *Ankrd31^-/-^* mice would have complicated interpretation of experimental outcomes. The first wave of spermatocytes reaches mid pachytene in 14 days old mice, at which point cellularities of testes in wild type and *Ankrd31^-/-^* mice begin to differ due to apoptosis of spermatogenic cells in the latter. Transcriptomes and histone H3K4me3 patterns are expected to greatly differ in meiotic and non-meiotic cell populations of testes. Therefore, altered relative proportions of these cell populations differentiate transcriptomes and histone H3K4me3 ChiPseq signal distributions in testes of wild-type and *Ankrd31^-/-^* mice that are older than 13-14days. Hence, we used 12 days old mice for these experiments.

Animals were used and maintained in accordance with the German Animal Welfare legislation (“Tierschutzgesetz”). All procedures pertaining to animal experiments were approved by the Governmental IACUC (“Landesdirektion Sachsen”) and overseen by the animal ethics committee of the Technische Universität Dresden. The licence numbers concerned with the present experiments with mice are T 2014-1 and TV A 8/2017.

### Generation of antibodies against fragments of ANKRD31, Histone H1t and REC114

Antibodies were raised against two ANKRD31 fragments (N-terminal fragment, 144 amino acids between Thr182 and Met325 residues, C-terminal fragment, 153 amino acids between Pro1470 and Arg1622 residues), full length Histone H1t, and two REC114 fragments (N-terminal, 130 amino acids between Met1 and Glu130 residues, C-terminal fragment, 129 amino acids between Arg131 and Asn259 residues). Coding sequences corresponding to these peptides were cloned into pDEST17 bacterial expression vector. Recombinant 6xHis-tagged proteins were expressed in *E. coli* strain, BL21 tRNA (ANKRD31) or BL21(DE3)pLysS (Histone H1t and REC114), and subsequently purified on Ni-Sepharose beads (Cat. no. 17-5318-01, Amersham, GE Healthcare). Purified proteins were used for immunization of rabbits and guinea pigs. ANKRD31 fragments coupled to NHS-Activated Sepharose 4 Fast Flow beads (Cat. no. 17-0906-01, Amersham, GE Healthcare) were used to affinity purify polyclonal antibodies following standard procedures.

### ANKRD31 immunoprecipitation and western blotting

For preparation of protein extracts from wild type and *Ankrd31* -deficient testes, testes of 12 days old juvenile mice were detunicated and homogenized in a lysis buffer (50 mM Tris-HCl pH 7.5, 150 mM NaCl, 0.5% Triton X-100, 1 mM MgCl_2_). Lysis buffer was supplemented with protease inhibitors and phosphatase inhibitors: 1 mM Phenylmethylsulfonyl Fluoride (PMSF); complete™ EDTA-free Protease Inhibitor Cocktail tablets (Roche, 11873580001); 0.5 mM Sodium orthovanadate; Phosphatase inhibitor cocktail 1 (Sigma, P2850) and Phosphatase inhibitor cocktail 2 (Sigma, P5726) were used at concentrations recommended by the manufacturers. Testis homogenates were lysed for 60 min at 4°C in the presence of benzonase (Merck Millipore) to digest DNA during lysis. Lysates were spun at 1000 g for 10 min. Supernatants were diluted two times with 50 mM Tris-HCl pH 7.5, 150 mM NaCl, mixed with 1 μg of rabbit anti-ANRKD31 C-terminal antibody and incubated for 2.5 h at 4° C. 1.5 mg of Dynabeads™ Protein A (Invitrogen) were added to the lysate-antibody mix and incubated for 4 h at 4°C. Beads were washed twice with washing buffer (50 mM Tris-HCl pH 7.2, 150 mM NaCl, 0.25% Triton X-100). Immunoprecipitated material was eluted from the beads by incubating the beads in 100 μl Laemmli sample buffer for 10 min at 70° C. The proteins from resulting elutions and input samples were separated on 4-15% TBX-acrylamide gradient gel (Bio-Rad) and blotted onto PVDF membrane (Sigma, P2938). Membranes were blocked for 1 h at room temperature using blocking solution (5% skimmed milk, 0.05% Tween 20, in TBS pH 7.6) and incubated overnight at 4°C with guinea pig anti-ANKRD31 N-terminal (1:1000) and mouse anti-GAPDH (1:1000) primary antibodies diluted in 0.05% Tween 20 in TBS pH 7.6 (TBS-T). Afterwards, horseradish peroxidase (HRP)-conjugated secondary antibodies (diluted in 2.5% skimmed milk in TBS-T) were applied for 1 h at room temperature. Detection of secondary antibodies was performed with Immobilon Western Chemiluminescent HRP Substrate (Millipore).

### Cytoplasmic and nuclear fractionation for PRDM9 detection

Juvenile mouse testes were homogenized in hypotonic buffer (10 mM Hepes, pH 8.0, 320 mM sucrose, 1 mM PMSF, 1x Complete protease inhibitor cocktail EDTA-free (Roche, Cat. Number 11873580001)) and phosphatase inhibitor 1x (Thermo Scientific, Halt Phosphatase Inhibitor Cocktail, Cat Number 78420) in a glass douncer. Testis cell suspensions were centrifuged at 1000g at 4°C for 10 min. Supernatants were collected and used as cytoplasmic fractions. Pellets were resuspended in RIPA buffer (50 mM Tris-HCl, pH 7.5, 150 mM NaCl, 1 mM EDTA, 1% NP-40, 0.5% Na-deoxycholate, 0.1% SDS, 1x Complete protease inhibitor EDTA-free (Roche)) and sonicated (4 cycles of 15s ON, 15s OFF, high power) on a Bioruptor Next-Gen sonicator (Diagenode). Suspensions were centrifuged at 16000 g, 4°C for 10min Then, supernatants were collected and used as nuclear fractions. Cytoplasmic and nuclear fractions (40 μg were separated on a 4-15% TBX-acrylamide gradient gel (Bio-Rad) and blotted onto nitrocellulose membranes. The membrane was blocked for 1 h at room temperature (1xTBS-T / 5% Milk) and cut according to expected sizes. The upper part was incubated over night at 4°C with affinity purified rabbit anti-PRDM9 (1:1000) (Grey et al., 2017) and guinea pig anti SYCP3 (1:3000) raised against mouse SYCP3 residues 24-44. Secondary antibodies were goat anti-rabbit IgG-HRP (1:5000) (1858415, Pierce) and goat anti-guinea pig IgG-HRP (1:5000) (706-035-148, Jackson Immuno Research). Blots were revealed with Super Signal West Pico Chemiluminescent Substrate, 34080, Thermo Scientific).

### Yeast two-hybrid (Y2H) assay

Yeast two-hybrid experiments were performed as described previously with minor modifications (Stanzione et al., 2016). Pairwise interactions were tested in the Y2HGold Yeast strain (Cat. no. 630498, Clontech). To transform Y2HGold with bait and prey vectors, yeasts were grown in 2xYPDA medium overnight at 30°C, 200 r.p.m. shaking. Afterwards, yeast cells were diluted to 0.4 optical density (measured at 600 nm) and incubated in 2xYPDA for 5 h at 30°C, 200 r.p.m. shaking. Cells were harvested, washed with water and resuspended in 2 ml of 100 mM lithium acetate (LiAc). 50 μl of this cell suspension was used for each transformation. Transformation mix included 1 μg of each vector (bait and prey), 60 μl of polyethylene glycol 50% (w/v in water), 9 μl of 1.0M LiAc, 12.5 μl of boiled single-strand DNA from salmon sperm (AM9680, Ambion), and water up to 90 μl in total. The transformation mix was incubated at 30°C for 30 min, and then at 42°C for 30 min for the heat shock. The transformation mix was removed following centrifugation at 1000 g for 10 min, and then cells were resuspended in water, and plated first on -Leu -Trp plates to allow selective growth of transformants. After 2 days of growth, transformants were plated both on -Leu -Trp and -Leu -Trp -Ade -His plates for 2-7 days to test for interactions. We followed the manufacturer’s instructions for media and plate preparation. The full length ANKRD31 protein activated the Y2H reporter system even in the absence of potential binding partner proteins from mice, which prevented the use of full length ANKRD31 in Y2H. To overcome this limitation, an array of fragments that cover the full length of ANKRD31 sequence were used in Y2H (Table S2). Self-activation tests showed that interactions could be tested along the entire length of ANKRD31 by using a combination of ANKRD31 prey and bait constructs.

### Immunofluorescence microscopy

#### Preparation of spermatocyte spreads

Preparation and immunostaining of nuclear surface spreads of spermatocytes was carried out according to earlier described protocols with minor modifications (Peters et al., 1997; Stanzione et al., 2016). Briefly, testis cell suspensions were prepared in PBS pH 7.4, then mixed with hypotonic extraction buffer in 1:1 ratio and incubated for 8 min at room temperature. After diluting the cell suspension five times in PBS pH 7.4, cell suspensions were centrifuged for 5 min at 1000 g, and cells were resuspended in the 1:2 mixture of PBS and 100mM sucrose solution. Cell suspensions were added to seven times higher volume droplets of filtered (0.2 μm) 1% paraformaldehyde (PFA), 0.15% Triton X-100, 1mM sodium borate pH 9.2 solution on diagnostic slides, and incubated for 60 min at room temperature in wet chambers. Nuclei were then dried for at least 1 h under fume-hood. Finally, the slides were washed in 0.4% Photo-Flo 200 (Kodak) and dried at room temperature.

#### Preparation of oocyte spreads

To prepare nuclear surface spread oocytes, two ovaries from each mouse were incubated in 20 μl hypotonic extraction buffer for 15 min (Hypotonic Extraction Buffer/HEB: 30 mM Tris-HCl, 17 mM Trisodium citrate dihydrate, 5 mM EDTA, 100 mM sucrose, 0.5 mM DTT, 0.5 mM PMSF, 1xProtease Inhibitor Cocktail). After incubation, HEB solution was removed and 16μl of 100 mM sucrose in 5mM sodium borate buffer pH 8.5 was added. Ovaries were punctured by two needles to release oocytes. Big pieces of tissue were removed. 9μl of 65 mM sucrose in 5 mM sodium borate buffer pH 8.5 was added to the cell suspension and incubated for 3 min. After mixing, 1.5μl of the cell suspension was added in a well containing 20μl of fixative (1% paraformaldehyde/50mM borate buffer pH:9.2/0.15% Triton-X100) on a glass slide. Cells were fixed for 45 min in humid chambers, then slides were air dried. Upon completion of drying slides were washed with 0.4% Photo-Flo 200 solution (Kodak, MFR # 1464510) for 5 min and afterwards they were rinsed with distilled water.

#### Immunofluorescence on gonad sections

To detect ANKRD31, testes were sectioned before fixation. Collected testes were immediately frozen in OCT (Sakura Finetek Europe) and sectioned (8 μm thick) on a cryostat. Slices of testes were allowed to dry on glass slides followed by fixation for 30 minutes at room temperature in 4% paraformaldehyde in phosphate buffer pH 7.4 followed by permeabilization in PBS/0.2% Triton-X100 for 10min. Slides were washed in PBS 3 times, and blocked in blocking buffer (5% BSA in PBS, 0.05% tween-20, 0.05% Triton X-100) before staining with anti-ANKRD31 and anti-SYCP3 antibodies, followed by DAPI staining. Anti-SYCP3 and DAPI staining served to facilitate exact staging of prophase in spermatogonial cells.

To detect apoptosis in testis or ovary sections, we sectioned testes both before and after fixation and ovaries only after fixation. To prepare sections of gonads after fixation, testes from adults and ovaries from newborn mice were fixed in 3.6% formaldehyde in PBS pH 7.4, 0.1% Triton X-100 at room temperature for 40 min (testes) or 20min (ovaries). After fixation testes/ovaries were washed 3 times in PBS pH 7.4 and placed in 30% sucrose overnight at 4°C. Fixed testes/ovaries were frozen on dry ice in OCT (Sakura Finetek Europe). 8 μm thick sections (testes) and 5 μm thick sections (ovaries) were cut and dried onto slides. For ovary sections, an additional step of permeabilization was performed by incubating the slides for 10 min in methanol and 1 min in acetone at -20 □C. The sections were washed in PBS pH 7.4 and immediately used for immunofluorescence staining. Anti-cleaved PARP staining (apoptosis marker) and histone H1t (post-mid pachytene stage marker) were detected by immunofluorescence in testes sections. DNA was counterstained by DAPI to facilitate staging of seminiferous tubules. Anti-cleaved PARP and GCNA1 (oocyte marker) (Enders and May, 1994)) were detected on oocyte sections. The numbers of cleaved PARP-positive and -negative oocytes were counted on every seventh section to determine the proportion of apoptotic oocytes.

To assess oocyte numbers in adult mice DDX4 was detected in paraffin-embedded sections of ovaries in young adults (6-7weeks old). Ovaries were dissected and fixed in 4% paraformaldehyde in 100 mM Sodium Phosphate buffer pH 7.4 overnight at 4°C. Afterwards, ovaries were washed 3 times in PBS pH 7.4, once with 70% ethanol and embedded in paraffin for sectioning at 5 μm thickness. Deparaffinization and rehydration of the sections was performed as follows: 2 x 5 min in xylene, 2 x 5 min in 100% ethanol, 5 min each in 95%, 85%, 70%, 50% ethanol, 2 x 5 min in water. Sections were subjected to heat-mediated antigen retrieval in 10 mM Sodium citrate, 0.05% Tween 20, pH 6.0 for 20min on boiling water bath. Sections were permeabilized in PBS with 0.2% Triton X-100 for 45 min at room temperature and processed for immunofluorescence staining immediately. DDX4-positive oocytes were counted on every seventh section of both ovaries in each female mouse.

#### Staining procedures

Previously described blocking and immunostaining procedures (Stanzione et al., 2016) were optimised for immunostaining with each combination of antibodies, details are available upon request.

#### Quantification of immunofluorescence signal levels and focus counts

Background corrected γH2AX signal was quantified in whole nucleus with a similar strategy as described earlier (Daniel et al., 2011). DSB repair foci were counted “manually” on matched exposure images of wild type and *Ankrd31M^-/-^* nuclear spreads of meiocytes. Focus numbers of pre-DSB recombinosome proteins (MEI4, REC114 and ANKRD31) and the co-localization of foci were quantified by Cell Profiler software as described previously for the analysis of MEI4, REC114 and IHO1 foci (Stanzione et al., 2016).

### Immunofluorescence staining combined with pseudoautosomal region (PAR) Fluorescence *in situ* hybridization (FISH)

After 10 min wash with Washing Buffer 1 (WB1, 0.4% Photo-Flo 200, 0.01% Triton X-100 in water), surface spreads were incubated overnight at room temperature with the primary antibody diluted in antibody dilution buffer (ADB) (10% goat serum, 3% bovine serum albumin [BSA], 0.05% Triton X-100 in phosphate-buffered saline [PBS]).

Then, slides were washed in 10 min WB1 and 10 min Washing Buffer 2 (WB2 0.4% Kodak Photo-Flo 200 in water), and incubated with the secondary antibody for 60 min in a pre-warmed humidified chamber at 37°C in the dark. Following further 10 min WB1, 10 min WB2 and 1 min PBS washes, slides were incubated in Hoechst 33258/PBS solution for at least 20 min in a room temperature humidified chamber. Air-drying slides for 10 min at room temperature in the dark, coverslips were mounted using ProLong^®^ Gold Antifade Mountant without DAPI (Molecular Probes- Life technologies cat. num. P36934). Images were captured using Leica CTR6000 Digital Inverted Microscope connected to a charge-coupled device camera and analyzed using the Leica software LAS-AF, for fluorescent microscopy.

#### Preparation of clone DNA and nick translation

BAC clone for PAR region (RP24500I4) were grown in standard Luria Bertani (L.B) medium at 32°C for a minimum of 16 h. DNA purification was obtained with Plasmid Maxi-Prep Kit (Qiagen). Determination of DNA concentration was made by both UV spectrophotometry at 260 nm and quantitative analysis on agarose gel. 1 μg of extracted BAC DNA was marked with fluorescent green-dUTP (Enzo Life Sciences, ref 02N32-050) using Nick Translation Kit (Abbott Molecular) according to the manufacturer’s instructions. For FISH experiment, marked probe was precipitated in EtOH 100%, 3 mM Sodium Acetate pH 5.2 and 0.1μg/μl mouse Cot-1 DNA (Invitrogen) at -80°C overnight. Resulting pellet was suspended in hybridization buffer (Enzo Life Sciences) according to the manufacturer’s instructions and denatured at 73°C for 5 min in a water bath.

#### PAR Fluorescence *in situ* hybridization (FISH)

Following immunofluorescence, spread slides were washed with fresh PBS at RT for 5 min, rinsed briefly in dH2O and dehydrated passing through an ethanol series and air-dried. After aging (65°C for 1 h), slides were denatured for 7 min in 70% formamide/2xSSC solution at 72°C and immediately dehydrated, passing it through -20°C cooled ethanol series and air-dried. FISH probe was applied to the slides for denaturation and hybridization steps in a humid chamber (75°C for 10 min and 37°C for at least 16 h, respectively). Following two washes with stringent wash buffer (4xSSC/0.2%Tween-20) at 55°C, slides were dehydrated through an ethanol series and air-dried. Nuclei were stained with Hoechst 33258/PBS solution for 20 min at RT and coverslips were mounted using Antifade Mountant without DAPI. Images were captured using Leica CTR6000 digital inverted microscope and analyzed using Leica software LAS-AF.

### Immunofluorescence staining combined with chromosome FISH

Testes from 14 days old mice (C57BL/6J) were dissected and spread nuclei of spermatocytes were prepared as described (Anderson et al., 1999). Briefly, a single-cell suspension of spermatogenic cells in 0.1 mM sucrose with protease inhibitors (Roche) was dropped on 1% paraformaldehyde-treated slides and allowed to settle for 3 h in a humidified box at 4°C. After brief washing with water and PBS, the spread nuclei were blocked with 5% goat serum in PBS (vol/vol), the cells were immunolabeled in a humid chamber at 4°C for 12 h with specific antibodies: anti-ANKRD31 N-terminal domain (diluted 1:400, rabbit antibody), anti-SYCP3 (1:50, mouse monoclonal antibody, Santa Cruz #74569) and anti-CENT (1:200, human – purified antibodies from human anti-centromere positive serum, Antibodies Incorporated #15-235). Secondary antibodies were used at 1:500 dilutions and incubated at 4°C for 90 min; goat antiRabbit IgG-AlexaFluor488 (MolecularProbes, A-11034), goat anti-Mouse IgG-AlexaFluor594 (MolecularProbes, A-11032), and goat anti-Human IgG-AlexaFluor647 (MolecularProbes, A-21445).

After the immunofluorescence staining, the slides were used for Fluorescence *in situ* hybridization (FISH) as described (Kauppi et al., 2011) with DNA FISH probes for mouse chromosomes 4, 9 and 13. Briefly, the slides were dehydrated, denatured and hybridized in a humid chamber at 37°C for 60 hours with one of three mouse chromosome painting probes with an orange emitting fluorochrome (Meta Systems probes): 1. XMP 4 orange (# D-1404-050-OR);

2. XMP 9 orange (# D-1409-050-OR); 3. XMP 13 orange (# D-1413-050-OR). After FISH, the slides were washed in stringency wash solution, drained and mounted in Vectashield-DAPI+ mounting medium.

The images were examined by Nikon Eclipse 400 microscope with a Plan Fluor objective 60x (MRH00601, Nikon) with a setup of five filter blocks: UV (Ex330-380/ Em400-420), FITC (Ex465-495/ Em505-520), ET Orange (Ex530-560/ Em580-600), Sp107/TR (Ex590-615/ Em615-645), CY5 (Ex620-660/ Em660-700). The images were captured using a DS-QiMc monochrome CCD camera (Nikon) and the NIS-Elements program (Nikon). The acquired images were processed with NIS-Elements program and saved in TIF for each channel.

### Diakinesis/Metaphase I chromosome spreading

Chromosome spreads of diakinesis/metaphase I stage spermatocytes were prepared as described in (Holloway et al., 2010). Briefly, testes were decapsulated and tubules were disrupted in hypotonic buffer (1% trisodium citrate in water). Large clumps were removed and the cell suspension was incubated in hypotonic buffer for 20min at room temperature. Cell suspension was centrifuged at 200g for 10min and supernatant was removed. Afterwards, cells were fixed in a methanol/acetic acid/chloroform (3:1:0.05 ratio) fixative, centrifuged and resuspended in ice-cold methanol/acetic acid solution (3:1 ratio). Fixed cells were dropped onto slides, dried quickly (in humid conditions), and stained with Hoechst 33342.

### XY chromosome painting in metaphase spreads

For sex chromosome painting, probes specific for X and Y chromosomes were used (XMP X Green, XMP Y Orange, MetaSystems). FISH protocol was performed according to manufacturer’s instructions.

### Staging of meiotic prophase

The first meiotic prophase can be subdivided into stages by a combination of three markers, SYCP3 (chromosome axis marker), SYCP1 (SC marker) and Histone H1t (post-mid pachytene marker in spermatocytes) (Table S1, for details see also (Stanzione et al., 2016). The stages that can be distinguished by these markers are listed briefly below, for a more detailed description see (Stanzione et al., 2016). Preleptotene stage corresponds to the premeiotic DNA replication in the cell cycle of germ cells. In a previous study (Stanzione et al., 2016), we used EdU labeling in combination with SYCP3 stain to define axis morphology that characterizes preleptotene spermatocytes. Based on this earlier study we define preleptotene as a stage where hazy/punctate staining pattern of SYCP3 is observed throughout the nucleus. The next stage, leptotene, is characterized by short stretches of axes and no SC. This is a stage where recombination is initiated in wild-type. The next stage is early zygotene, which is characterized by long, yet still fragmented, axis stretches. SC is also detected in this stage in SC proficient genotypes. During late zygotene, axes of all chromosomes are fully formed but SCs are incomplete. Cells enter pachytene when SC formation is completed in wild-type. All chromosomes are fully synapsed in pachytene oocytes. In contrast, only autosomes synapse fully and heterologous sex chromosomes synapse only in their PARs in spermatocytes. We also define a zygotene-to-pachytene transition stage for spermatocytes, where up to one autosome pair has not finished synapsis and/or sex chromosomes were still unsynapsed or about to synapse in wild-type. Sex chromosome axes, in particular X chromosome axes, are long and stretched out in this stage. Histone H1t staining is used to sub-stage pachytene. Histone H1t is absent or weak in early pachytene. Histone H1t levels are intermediate and high in mid and late pachytene, respectively. Pachytene is followed by the diplotene stage, during which axes desynapse and the SC becomes fragmented. Histone H1t levels are high in this stage in spermatocytes. In oocytes the same stages exist but histone H1t cannot be used as a staging marker. Instead the developmental time of fetuses can be used to aid staging. Most oocytes are in zygotene and mid-pachytene in foetuses 16 and 18 days postcoitum. Most oocytes are in pachytene/diplotene in newborn mice.

Staging of prophase beyond early zygotene is not straightforward in mutants that have SC defects. This is because fully formed axis characterizes stages between late zygotene and early diplotene, and SC cannot be used as a reliable marker in cells where SC formation is incomplete. We used histone H1t levels to aid staging in spermatocytes. Given the delayed/defective SC formation in *Ankrd31M^-/-^* mice the staging of late zygotene-like cells is uncertain. Hence, *Ankrd31M^-/-^* spermatocytes that have fully formed axis, incomplete SC, and low histone H1t levels could correspond to late zygotene or early pachytene wild type spermatocytes. Given the PAR synapsis defect in *Ankrd31M^-/-^* spermatocytes we defined zygotene-to-pachytene transition as a stage where up to one autosome pair is not fully synapsed in *Ankrd31M^-/-^* spermatocytes. However, we note that some of these cells might be early pachytene cells with defective synapsis. Pachytene was defined as a stage where all autosomes are fully synapsed but sex chromosomes may or may not be synapsed in *Ankrd31M^-/-^* spermatocytes.

### Staging of mouse seminiferous tubule cross sections

Spermatogenic cells that are located in the same section of a seminiferous tubule initiate and execute meiosis on a coordinated manner in mice. Meiotic entry occurs in spermatogenic cell layers at the perimeter of seminiferous tubules. Upon progression in meiosis, spermatogenic cells move towards the lumen of tubules. Concurrently, mitotic proliferation generates a new layer of spermatogenic cells for a new wave of meiosis at the perimeter of each seminiferous tubule. The combination of repeated meiosis entry and spermatogenic cells migration to the lumen generates the so called epithelial cycle of seminiferous tubules. In mice, the seminiferous epithelial cycle has 12 well-defined stages (I-XII) which are characterized by distinct associations of premeiotic, meiotic and post meiotic spermatogenic cell layers across crossections of seminiferous tubules (Ahmed and de Rooj, 2009). As spermatogenesis progresses, each portion of seminiferous tubules transits from stage I to XII, and then to stage I to start a new cycle. Nuclear morphology and chromatin condensation patterns differ in distinct stages of the spermatogenic process, hence detection of chromatin by DAPI was used to identify spermatogenic cell associations that define distinct stages of the seminiferous epithelial cycle. In some cases we also immunostained SYCP3 (axis) or histone-H1t (marker of spermatocytes after mid pachytene) to aid staging of the epithelial cycle. Staging without molecular markers of chromosome axis and SC suggested that stage X of epithelial cycle is characterized by the combination of elongating spermatids in the lumen, a mixture of late pachytene and diplotene cells in an intermediate cell layer, and spermatocytes that transit from leptotene to zygotene in the basal layer of seminiferous tubules (Ahmed and de Rooij, 2009). Immunostaining of axis, unsynapsed axis and/or SC markers (Figure 1C, S4E and our unpublished observations) suggested that elongating spermatids are present mostly in combination with diplotene and zygotene cells in our strain background. Hence we labeled basal and intermediate cell layers as zygotene and diplotene in Figure 1C and S4E.

### Immunofluorescence-based assessment of recombination kinetics

#### SC formation kinetics

To assess SC formation we quantified SC formation in prophase stages that were identified based on axis morphology (Figure 3D and S5B). We focused on pre-mid pachytene stages in spermatocytes because autosomal SC defects cause spermatocyte elimination in mid pachytene (Burgoyne et al., 2009; Mahadevaiah et al., 2008), which renders later stages of prophase unreliable for the quantification of autosomal SC defects in mutants with compromised synapsis. Hence, we quantified SC formation in juvenile mice before most spermatocytes reach mid pachytene, or we examined histone H1t (marker of post-mid pachytene stages) negative spermatocytes from adult (Figure 3D). Quantification of SC formation has limitations. Reduction in the number of cells that completed SC formation between autosomes can reflect either a kinetic delay of SC formation and/or a terminal failure of SC formation in a subpopulation of early pachytene spermatocytes. Hence, it was not possible to judge if there was a significant terminal failure in SC formation or only a kinetic delay in *Ankrd31^-/-^* spermatocytes.

#### DSB repair focus kinetics

To assess recombination kinetics we used immunostaining of DSB repair proteins that accumulate on the processed single-stranded DNA ends that result from DSBs (RPA, DMC1 and RAD51). This type of analysis provides information about the steady state level of recombination intermediates that are marked by the respective proteins. Thus the number of foci could vary with both the formation kinetics and the turnover of intermediates. Hence, focus numbers do not directly reflect DSB numbers. Nonetheless, the numbers of RAD51/DMC1 foci are thought to reflect the numbers of unrepaired DNA ends that are available for homology search in the context of unsynapsed chromosome axes. RPA focus counts are thought to reflect the number of all the single-stranded DNA ends that are participating in recombination both in the context of unsynapsed and synapsed chromosome axes.

### Gene expression analysis

In order to test the effect of *Ankrd31* deficiency in mouse testicular transcriptome total RNAs from *Ankrd31^+/+^* and *Ankrd31^-/-^*juvenile testes (12 days old) were extracted. RNA was extracted using RNeasy Mini (Qiagen, Cat No./ID: 74104) according to manufacturer’s instructions. Quantification and quality control of RNA was performed using Agilent 2100 Bioanalyzer. mRNA was isolated from 300 ng total RNA by poly-dT enrichment followed by strand specific RNA-Seq library preparation (Ultra II Directional RNA Library Prep, NEB) following the manufacturer’s instructions. Libraries were equimolarly pooled and subjected to 76 bp single end sequencing on a NextSeq 500 and Hiseq 2500 sequencer (Illumina) resulting in on average 29 Mio reads/sample (Bray et al., 2016; Kim et al., 2013).

To screen for genes that are preferentially expressed in gonads, we profiled transcriptomes of embryonic female gonads, adult testes and an array of 17 somatic tissues from adult mice by RNA-sequencing of total RNAs. RNeasy Mini kit (Qiagen, Cat No./ID: 74104) was used according to manufacturer’s instructions to purify total RNA from testes of adult mice and ovaries from fetal mice (C57/BL6xDBA/2 background). RNA was purified from ovaries that were pooled from 34-44 fetuses/newborn mice at six developmental timepoints, 11.5, 12.5, 14.5, 16.5, 18.5 and 20.5 days post coitum. Total RNA samples from mouse somatic tissues were purchased via Ambion (liver, brain, thymus, heart, lung, spleen and kidney, Cat#7800) and Zyagen (mammary gland, pancreas, placenta, salivary gland, skeletal muscle, skin, small intestine, spinal cord, tongue and uterus, Cat#MR-010). Total RNA from the listed 17 somatic tissues were mixed in equal proportions to create a somatic RNA mix, and total RNAs from testes were mixed with this somatic RNA mix in 1:17 ratio to create a testis+somatic RNA mix. mRNAs were isolated from 300 ng total RNA by poly-dT enrichment followed by strand specific RNA-Seq library preparation (Ultra II Directional RNA Library Prep, NEB) following the manufacturer’s instructions. Libraries were subjected to 75 bp single end sequencing on a NextSeq 500 (Illumina).

### ChIP experiments

DMC1 ChIP-seq was performed as described in (Grey et al., 2017). Two testis from *Ankrd31^+/+^* and *Ankrd31^-/-^* from 6 weeks old mice were used for each replicate. Sequencing was performed on HiSeq 2500 Rapidmode 2×50b.

H3K4me3 ChIP-seq was done as described in (Diagouraga et al., 2018). Four testis from *Ankrd31^+/+^* and *Ankrd31^-/-^* from 12dpp mice were used for each replicate. Sequencing was performed on HiSeq 2500 Rapidmode 1×75b.

### ChIP-seq data computational analysis

#### Read alignment

After quality control, H3K4me3 ChIP-seq and DMC1 ChIP-SSDS (Single Strand DNA Sequencing) reads were mapped to the UCSC mouse genome assembly build GRCm38/mm10. Bowtie2 (with default parameters) was used to map H3K4me3 ChIP-seq reads, while the previously published method (Khil et al., 2012) was used for DMC1 ChIP-SSDS reads (*i.e.* the BWA modified algorithm and a customized script, which were specifically developed to align and recover ssDNA fragments). A filtering step was then performed on aligned reads for all ChIP-seq experiments to keep non-duplicated and high-quality mapped reads with no more than one mismatch per read. When analyzing specifically the DMC1 and H3K4me3 signals covering the pseudo-autosomal region (PAR) of sex chromosome (Fig. 6F-G, Table S7), we used a read mapping performed over a customized genome which was depleted for the PAR from assembled chrY (*i.e.* only chrY:0-90745844 was included in this customized genome). This method ensured that all reads originated from either chrX- or chrY-PAR region were mapped over a unique reference sequence, preventing multimapping and thus misquantification over this region.

#### Identifying meiotic hotspots

To identify meiotic hotspots from biologically replicated samples in DMC1 ChIP-SSDS, we used the Irreproducible Discovery Rate (IDR) methodology, as previously described (Diagouraga et al., 2018). This method was developed for ChIP-seq analysis and extensively used by the ENCODE and modENCODE projects (Landt et al., 2012). The framework developed by Qunhua Li and Peter Bickel’s group (http://sites.google.com/site/anshulkundaje/projects/idr) was followed. Briefly, this method allows testing the reproducibility within and between replicates by using the IDR statistics. Following their pipeline, peak calling was performed using MACS version 2.0.10 with relaxed conditions (--pvalue=0.1 --bw1000 --nomodel --shift400) on each of the two replicates, the pooled dataset, and on pseudo-replicates that were artificially generated by randomly sampling half of the reads twice for each replicate and the pooled dataset. Then IDR analyses were performed and reproducibility was checked. Final peak sets were built by selecting the top N peaks from pooled datasets (ranked by increasing p values), with N defined the highest value between *n_1_* (the number of overlapping peaks with an IDR below 0.01, when comparing pseudo replicates from pooled datasets) and *n_2_* (the number of overlapping peaks with an IDR below 0.05, when comparing the true replicates), as recommended for the mouse genome.

#### Signal normalization and quantitative analysis

All read distributions and signal intensities presented in this work were then calculated after pooling reads from both replicates. DMC1 ChIP-SSDS peaks were re-centered and read enrichment was normalized to the local background, as previously described (Brick et al., 2012). DMC1 ChIP-SSDS read enrichments could not be normalized between *Ankrd3^+/+^* and *Ankrd31^-/-^* mice. Indeed, the previously reported method to normalize such data (that is to consider that the total DMC1 signal should be the same between genotypes□; (Davies et al., 2016; Diagouraga et al., 2018) cannot apply to our study as the DMC1 dynamics is altered in the mutant (see Figure 4). Thus, only relative levels between regions or hotspots should be compared between different genotypes. SSDS counts from DMC1 ChIP are dependent on DMC1 half-life on ssDNA and are normalized to library size hence they can be only used to assess distribution but not the absolute numbers of single-stranded-DNA-containing recombination intermediates. The H3K4me3 signal recovered from young mice has been normalized between *Ankrd3^+/+^* and *Ankrd31^-/-^* mice assuming, as in previous studies (Davies et al., 2016; Diagouraga et al., 2018), that the signal of a PRDM9-independent set of H3K4me3 peaks should remain unchanged in testes of fertile mice at the same meiotic stage. Normalization factors were thus calculated based on the enrichment found at a set of 29 promoters of protein-coding genes (*Mlh1, Pms2, Mnd1, Dmc1, Rad21l, Mei4, Stra8, Ctcfl, Mei1, Puf60, Eef2, Rpl38, Leng8, Setx, Eif3f, Rpl37, Psmd4, Heatr3, Chmp2a, Sycp1, Sycp2, Sycp3, Morc2b, Zfp541, Spo11, Mdh1b, Rec8, Msh4, Psmc3ip*), suggested by Davies et al (2016). H3K4me3 signal presented in the present study were thus normalized by library size (Read per million of mapped reads; RPM) then to promoters to reach the level of *Ankrd31^+/+^.*

#### Determination of overlapping peaks

DSB hotspots identified in *Ankrd31^-/-^* mice were assigned to B6 or default hotspots on the basis of overlapping peak centers ±200bp. As reference for B6 hotspots we used previously published DSB maps (Diagouraga et al., 2018; Grey et al., 2017; Smagulova et al., 2016) and the DSB map of *Ankrd31^+/+^* mice used in this study. As a reference for default DSB sites, we used the map of the *Prdm9^-/-^* mice (Brick et al., 2012). When necessary, DSB hotspot coordinates were converted from GRCm37/mm9 to GRCm38/mm10 using the liftOver tool (http://genome.ucsc.edu/cgi-bin/hgLiftOver). Thus, 94% of peaks could be assigned to a known DSB hotspot, whereas a small fraction (6%) of DSB hotspots could not be assigned to any of the reference hotspots and was defined as new (Fig. 6B). These new hotspots have in average a weak SSDS signal (Fig. 6C) and could be PRDM9 dependent or independent.

### Statistical analysis

Statistical analysis of cytological observations was done by GrapPad Prism 7. All other statistical tests were done using R version 3.3.3, if not otherwise stated. All tests and p-values are provided in the corresponding legends and/or figures.

### ANKRD31 family collection and protein sequence analysis

A Hidden Markov Model (HMM) search with the mouse ANKRD31 protein in the PFAM database (v. 31.0, March 2017, (Finn et al., 2016)) detected two ankyrin repeat regions, with 3 copies each (amino acids 467-568 and 1167-1279) with highly significant E-values (1.6e-35). To search for orthologs, we performed a NCBI blastp search within the complete set of UniProt reference proteomes or the NCBI non-redundant protein database, restricting the query to the region c-terminal of the last ankyrin repeat (amino acids 1280-1856) (Sayers et al., 2011). The Ankrd31 protein family is well conserved within vertebrates and orthologs can be collected in reciprocal blasts by applying highly significant e-values (1e-10). For a multiple alignment, sequences were derived either from the UniProt database, as for the Ankrd31 orthologs of Homo sapiens (sp|Q8N7Z5|ANR31_HUMAN), Ornithorhynchus anatinus (tr|F7FXW5|F7FXW5_ORNAN), and Xenopus tropicalis (tr|F6ZDM1 |F6ZDM1_XENTR), or from the NCBI protein database, as for Canis lupus familiaris (ref|XP_022272501.1|), Mus musculus (ref|XP_006517860.1|), Columba livia (ref|XP_021147901.11), Chrysemys picta bellii (ref|XP_023960269.11), Struthio camelus australis (ref|XP_009686903.1|), Callorhinchus milii (ref|XP_007903536.1|), and Takifugu rubripes (ref|XP_011613543.1|). Multiple alignment was performed with MAFFT (-linsi v7.313, (Katoh and Toh, 2008)), and visualized with ClustalX (v2.1,(Thompson et al., 1997)). 5 conserved regions were identified. The first ankyrin repeat region is missing in bony fish, but present in shark, whereas the second ankyrin repeat region is highly conserved in all vertebrate orthologs. The third conserved region CR3 includes a predicted coiled coil interaction domain (in mouse ANKRD31 from 1340 to 1374 (Lupas et al., 1991)). For the conserved regions CR4 (1701-1787) and CR5 (1811-1857) no function could be assigned.

## Supplemental Information titles and legends

**Figure S1 related to Figure 1.**
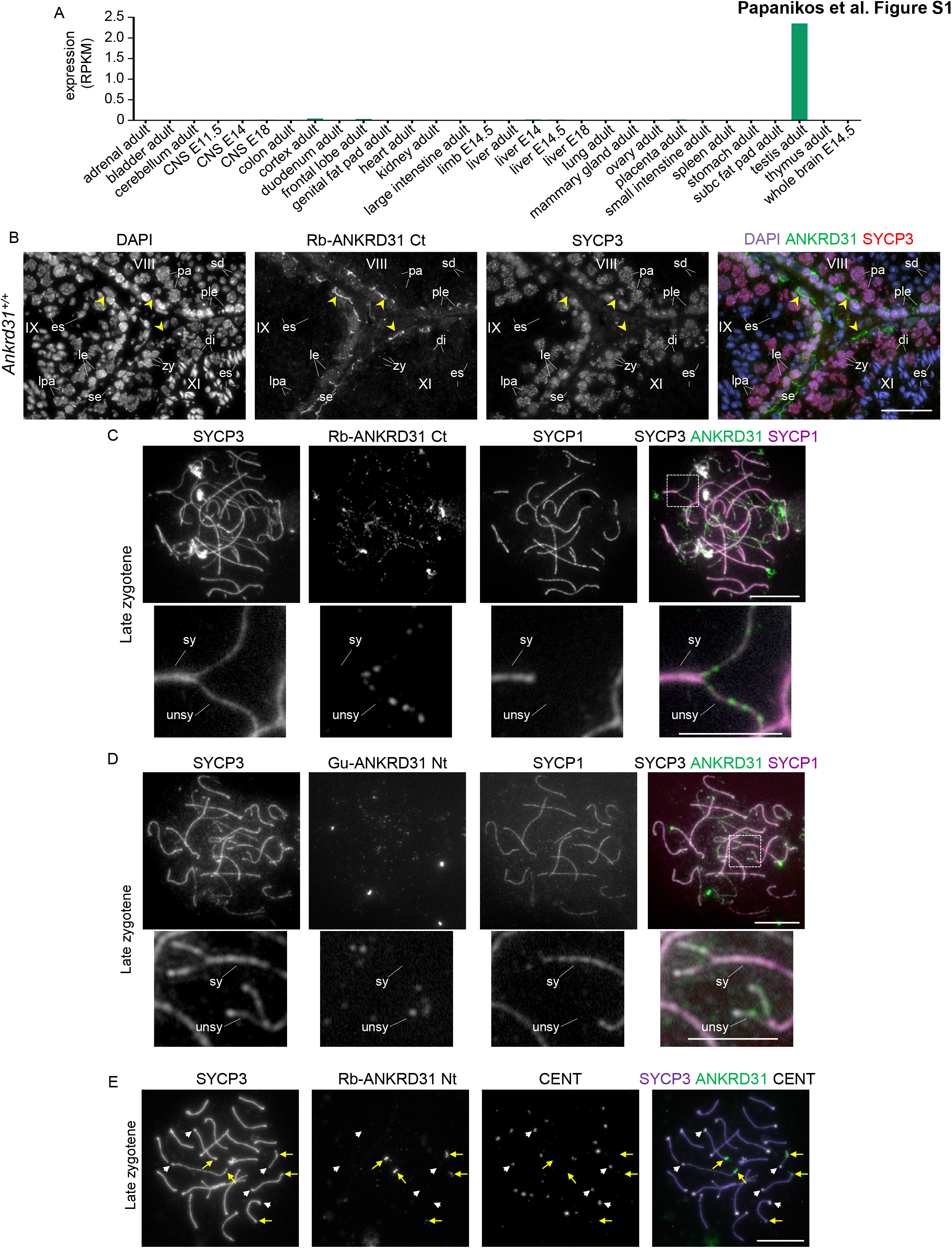
(**A**) *Ankrd31* transcript levels are shown as reads per kilo base million (RPKM) in RNAs of indicated tissues from the ENCODE project (data source, GEO accession: GSE36025; http://www.ncbi.nlm.nih.gov/gene/625662/?report=expression). (**B**) DNA was detected by DAPI on cross sections of adult mouse testes. Immunofluorescence was used to detect ANKRD31 and a marker of meiotic chromosome axis, SYCP3. Antibodies raised against a C-terminal fragment of ANKRD31 in rabbit (Rb-ANKRD31 Ct) were used. Seminiferous tubules at stages VIII, IX and XI are shown. Sertoli cells (se), preleptotene (ple), leptotene (le), zygotene (zy), late pachytene (lpa) and diplotene (di) spermatocytes, round (sd) and elongating (es) spermatids are marked. The image shows a strong extranuclear signal (yellow arrowhead) that was detected by the Rb-ANKRD31 Ct antibodies between mitotic germ cells or preleptotene spermatocytes and more advanced spermatocytes. We consider this signal non-specific as the guinea pig anti-ANKRD31 antibodies did not produce similar pattern, see also Figure 1C. (**C, D**) Immunofluorescence staining of the indicated proteins is shown on nuclear surface spread spermatocytes of adult mice in late zygotene. SYCP1 is a marker of synapsed chromosomal regions. ANKRD31 was detected by Rb-ANKRD31 Ct (**C**) or antibodies raised against an N-terminal fragment of ANKRD31 in guinea pig (Gu-ANKRD31 Nt, **D**). Enlarged insets of top panels are shown in the bottom panels. Synapsed (sy) and unsynapsed (unsy) chromosome axes are indicated. (**E**) SYCP3, ANKRD31 and centromeric proteins (CENT) were detected by immunofluorescence. A late zygotene spermatocyte is shown. Yellow arrows point at ANKRD31 aggregates at the non-centromeric ends of five chromosomes. The centromeric ends of the same chromosomes are marked by white arrowheads. Bars, 50μm (**B**), 10μm (**C, D, E**), 5μm (**C** and **D** in enlarged insets).

**Figure S2 related to Figure 1 and 2.**
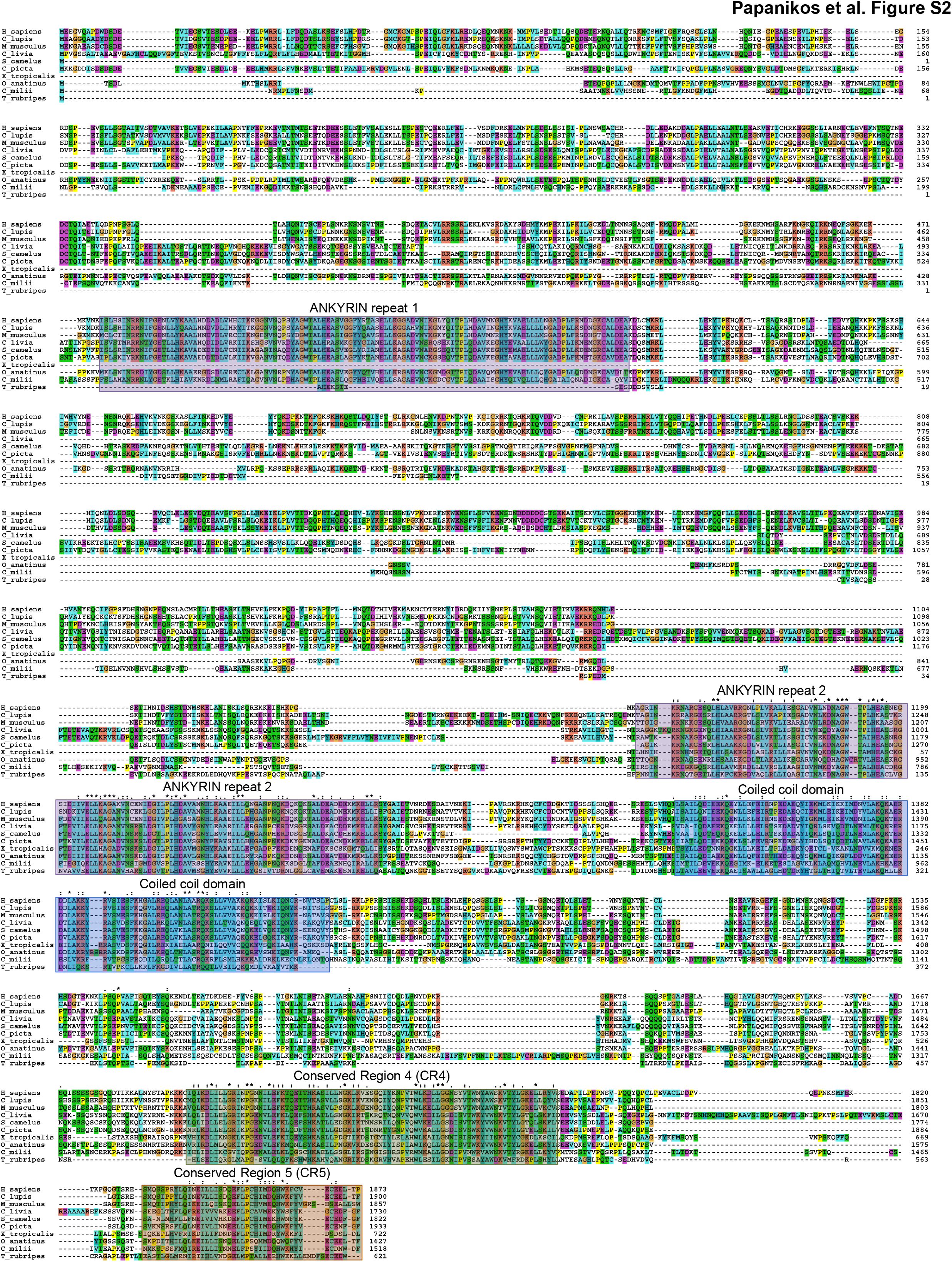
Multiple alignment of ANKRD31 protein sequences from various vertebrate taxa. Sequences are aligned from human (H_sapiens), dog (C_lupus), house mouse (M_musculus), platypus (O_anatinus), rock dove (C_livia), common ostrich (S_camelus), painted turtle (C_picta), western clawed frog (X_tropicalis), australian ghost shark (C_milii), japanese puffer fish (T_rubripes). Default ClustalX colour scheme was used. An “μ” (asterisk) indicates positions which have a single, fully conserved residue. A “:” (colon) indicates conservation between groups of strongly similar properties, scoring > 0.5 in the Gonnet PAM 250 matrix. A “.” (period) indicates conservation between groups of weakly similar properties, scoring =< 0.5 in the Gonnet PAM 250 matrix (Larkin et al., 2007). Boxes indicate five evolutionarily conserved regions of ANKRD31. We note that the more N-terminally located Ankyrin repeat domain is less conserved as this domain is missing in the ANKRD31 of bony fish.

**Figure S3 related to Figure 2.**
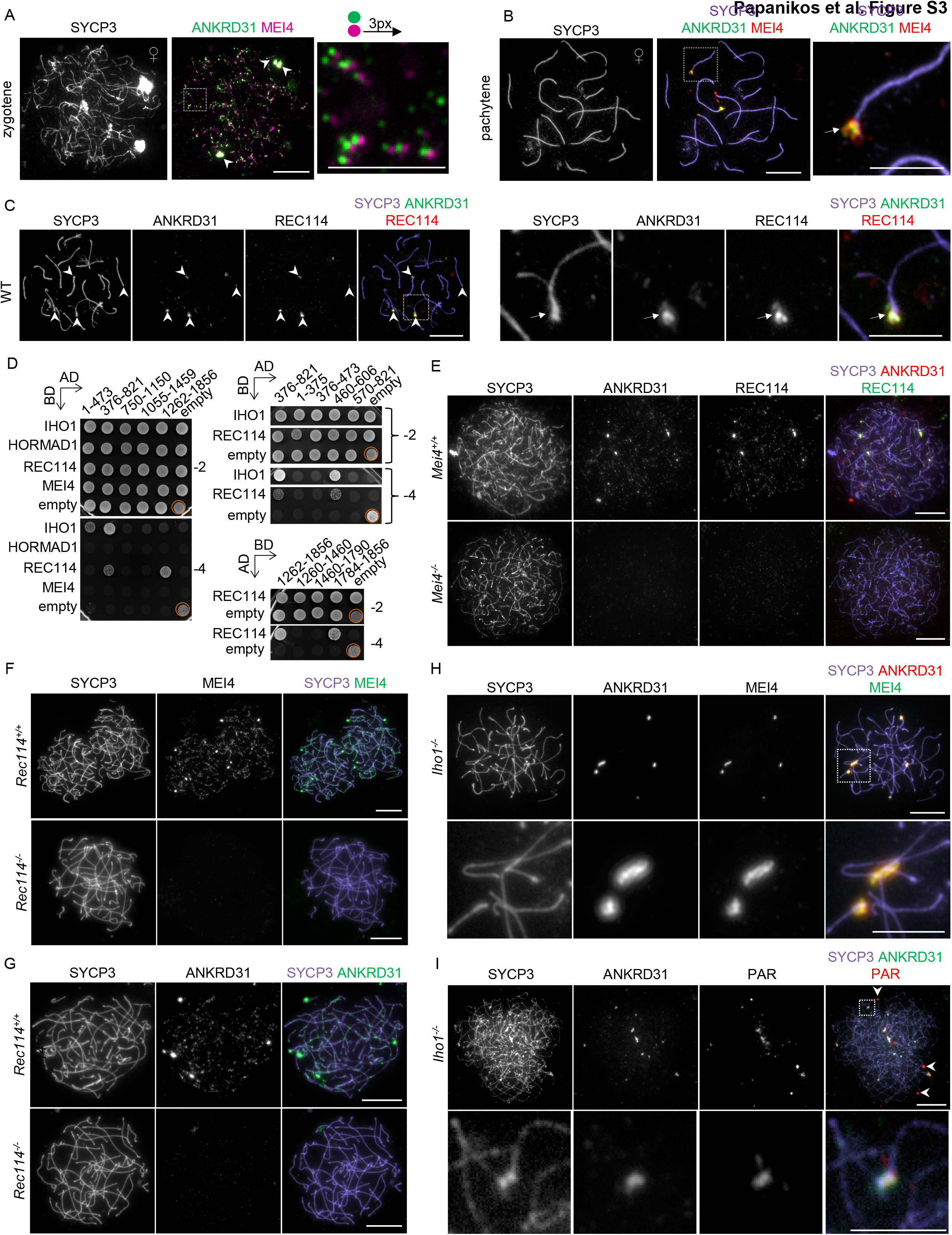
Indicated proteins were detected by immunofluorescence in nuclear spread oocytes (**A**, **B**) or spermatocytes (**C, E-I**). (**A**) A zygotene oocyte is shown from a fetus at 16 days post coitum (dpc). MEI4 signal is shifted to the right with three pixels in the enlarged insets to facilitate detection of overlap with focal ANKRD31 signal. Arrowheads mark large co-aggregates of ANKRD31 and MEI4. The median number of aggregates was four in late zygotene oocytes, n=24. (**B**) Pachytene oocyte is shown from an 18 dpc fetus. Enlarged inset shows a persisting co-aggregate of MEI4 and ANKRD31 (white arrow) on a synapsed chromosomal end. 25 of 44 (57%) of early-mid pachytene oocytes contained ANKRD31 aggregates, and the median number of aggregates was two. (**C**) Co-aggregates of ANKRD31 and REC114 are shown (marked by arrowheads) at the ends of three autosomes and the paired PARs of sex chromosomes in a spermatocyte at zygotene-to-pachytene transition. The four images on the right show an enlarged version of the boxed area in the left panel and arrow marks PAR. (**D**) Yeast two-hybrid assays testing interactions of indicated proteins with fragments of ANKRD31 (amino acid positions of fragment ends are indicated); AD – Gal4 activation domain, BD – Gal4 binding domain. 10 μl of yeast cell suspensions of optical density 0.5 were plated onto dropout plates – Leu, -Trp (-2) and -Leu, -Trp, -His, -Ade (-4). The plates were imaged after 2 days of growth. Orange circles show positive control Gal4-BD-HORMAD1/Gal4-AD-IHO1. IHO1 and REC114 were tested for interaction with ANKRD31 fragments that covered the first 821 amino acids of ANKRD31 (upper right panel) in the same experiment, therefore only one negative control is shown for this experiment (indicated by brackets). (**E-G**) show spermatocytes at stages that are equivalent to late zygotene and early pachytene in wild-type as judged by fully formed chromosome axes. (**H**) Enlarged insets (bottom panel) show co-aggregates of ANKRD31 and MEI4. (**I**) SYCP3 and ANKRD31 were detected by immunofluorescence, and PARs were detected by FISH in surface spread spermatocytes. Three *IhoT^-/-^* spermatocytes (upper panel) are shown at a stage that corresponds to late zygotene in wild-type based on axis morphology. Unspecific PAR signal that is not associated with chromosome axes is marked (white arrowhead) in the overlay image. Enlarged insets in lower panel show ANKRD31 aggregates that associate with two chromosomal ends with PAR FISH signal. Bars are 10μm and 5μm in low resolution images and enlarged insets.

**Figure S4 related to Figure 3.**
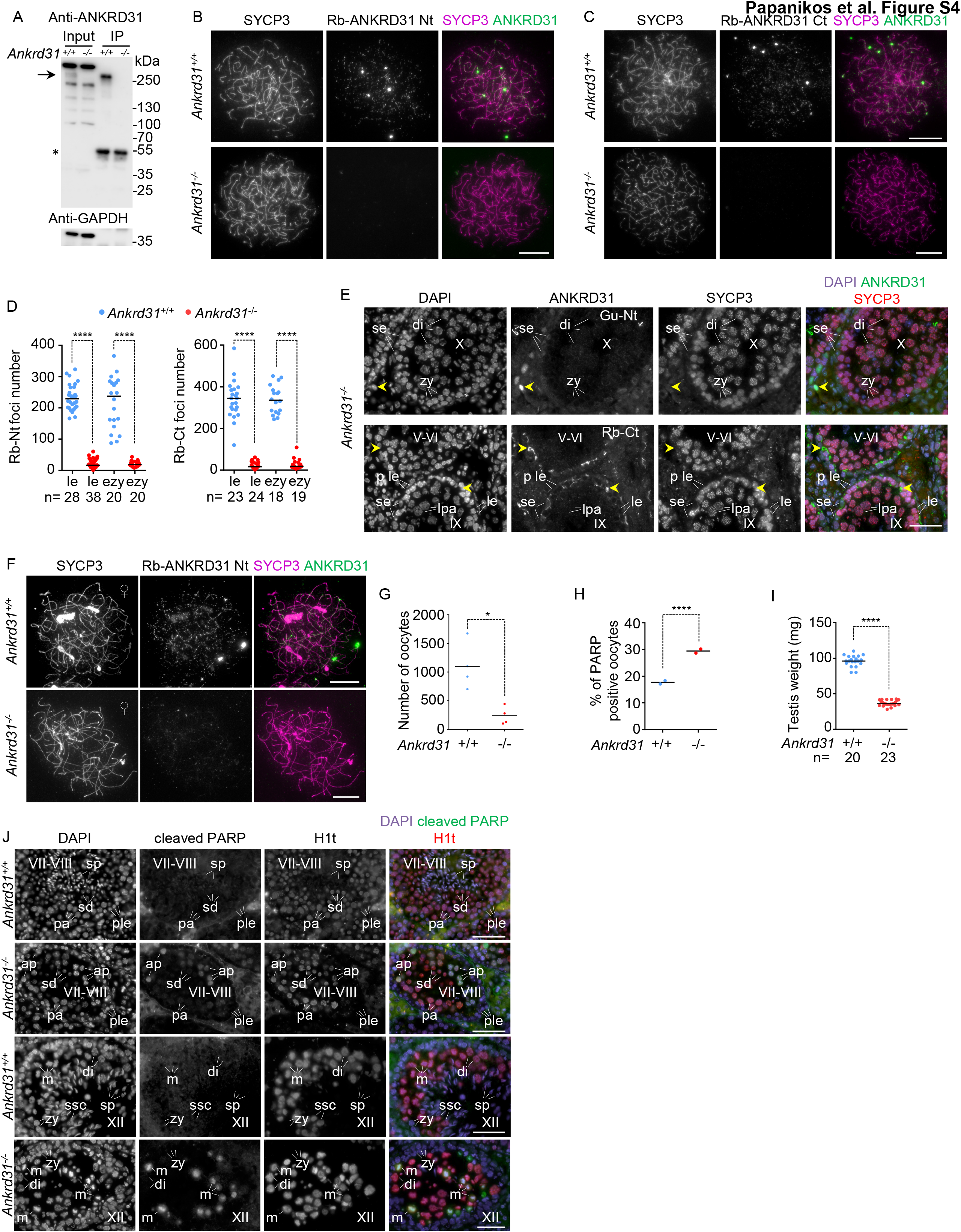
**(A)** ANKRD31 was immunoprecipitated from testes extracts of 12 days old *Ankrd31^+/+^* and *^-/-^* mice by rabbit antibodies raised against a C-terminal fragment of ANKRD31 (fragment: Pro1470 to Arg1622). Immunoblot analysis of immunoprecipitation input samples and immunoprecipitates (IP) is shown. Upper image shows detection of proteins on the blot membrane by guinea pig antibodies raised against an N-terminal fragment of ANKRD31 (fragment: Thr182 to Met325). Arrow marks presumed band of ANKRD31, which is present only in wild-type samples. Asterisk marks heavy chain of antibodies used for immunoprecipitations. Molecular weight marker positions are indicated. Anti-GAPDH (bottom panel) was used to control for loading in input samples. (**B, C, E, F, J**) Indicated proteins were detected in nuclear surface spreads of spermatocytes in **B**, **C** or spread oocytes in **F**, or cross sections of testes from adult mice of indicated genotypes in **E** and **J**. (**D)** Quantification of focal staining patterns by rabbit antibodies that recognise either an N-terminal (Rb-Nt) or a C-terminal (Rb-Ct) fragment of ANKRD31. Numbers of foci are shown in leptotene (le) and early zygotene (ezy) stages. Medians and number (n) of counted cells are indicated. Mann-Whitney U test indicate significantly lower focus numbers in *Ankrd31*^-/-^than *Ankrd31^+/+^* spermatocytes, **** corresponds to P<0.0001. (**E, J**) DNA was detected by DAPI. Stages of seminiferous tubules are indicated. (**E**) ANKRD31 was detected by either guinea pig antibodies raised against an N-terminal fragment of ANKRD31 (Gu-Nt, top panel), or rabbit antibodies raised against a C-terminal fragment of ANKRD31 (Rb-Ct, bottom panel). To compare ANKRD31 signal in testis sections of *Ankrd31^+/+^* and *Ankrd31^-/-^* see Figure 1C and Figure S1B. ANKRD31 images in top and bottom panels correspond to matched exposure images of ANKRD31 in Figure 1C and Figure S1B, respectively. Yellow arrowheads mark strong nuclear staining in an intertubular cell (upper panel) and strong extranuclear staining in the vicinity of the basal layer of spermatogenic cells (lower panel). These staining patterns are not shared by the Gu-Nt and Rb-Ct antibodies, hence these signals are not considered to represent ANKRD31 protein. We note that nuclear staining of ANKRD31 in zygotene (Figure 1C) or leptotene (Figure 1C and Figure S1B) spermatocytes of *Ankrd31^+/+^* contrasts the “negative” ANKRD31 staining in the nucleus of zygotene (zy, top panel) and leptotene (le, bottom panel) spermatocytes of *Ankrd31^-/-^* mice. Sertoli cells (se), leptotene (le), zygotene (zy), late pachytene (lpa) and diplotene (di) spermatocytes are marked. (**G**) Quantification of oocyte numbers in ovaries of 6-7 weeks old females. Ovaries were sectioned and oocytes were counted in every 7^th^ section. Data points show the sum of counts from the two ovaries of each mouse. Averages are indicated. Two tailed t test with Welch correction calculated statistical significance at 0.05>P>0.01 (*). (**H**) Graph shows fractions of oocytes that are apoptotic, as indicated by positive staining for cleaved PARP, in ovaries of newborn mice of indicated genotypes. Each data point represents fractions of oocytes in one mouse. Averages are shown. Chi square statistics calculated significance at P<0.00001 (****). (**I**) Quantification of testis weight in adult *Ankrd31^+/+^* and *Ankrd31^-/-^* mice. Each data point represents the sum of weights of two testes from one mouse. Medians are shown, Mann-Whitney U test calculated significance at P<0.0001 (****). (**J**) Apoptosis was detected by cleaved PARP staining in histological sections of testes in indicated genotypes. Histone H1t (H1t), which is a marker of post-mid pachytene stages in spermatocytes, was detected to aid staging of seminiferous tubules. Seminiferous tubules at stage VII-VIII (upper two panels) and XII (lower two panels) are shown. Preleptotene (ple), pachytene (pa) and apoptotic pachytene (ap) spermatocytes and round spermatids (sd) and sperm (sp) are marked in stages VII-VIII. Zygotene (zy), diplotene (di), metaphase (m) and secondary (ssc) spermatocytes and sperm (sp) are marked in stage XII. Bars, 50μm in **E** and **J**, 10μm in **B, C, F**.

**Figure S5 related to Figure 3 and 4.**
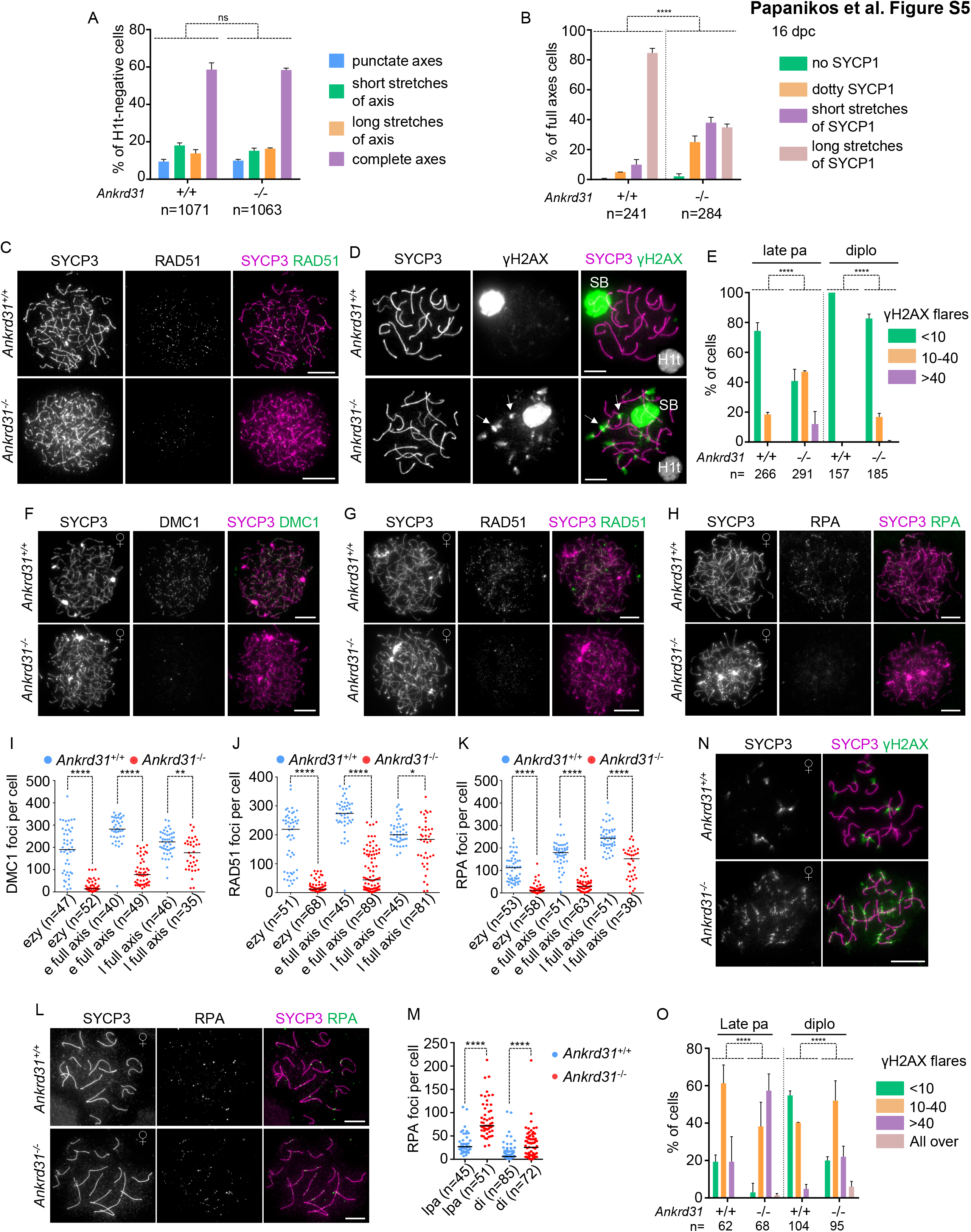
**(A**) Quantification of testicular cell populations with distinct chromosome axis morphologies. SYCP3 (axis marker) and histone H1t were detected in testicular cells from adult mice. Graph shows the proportion of pre-mid pachytene (H1t-negative) spermatocytes that have focal SYCP3 staining (punctate axes, corresponds to preleptotene), short stretches of axes (corresponds to leptotene), long stretches of axis (corresponds to early zygotene) or fully formed axes on all chromosomes (complete axes, corresponds to late zygotene and early pachytene). Standard deviation and weighted averages of percentages are shown from three experiments. Chi Square statistics indicated no significant (ns) difference. (**B**) Quantification of synaptonemal complex formation based on SYCP1 staining in oocytes of 16 dpc fetuses of indicated genotypes. Categories of SYCP1 staining patterns are indicated in oocytes with fully formed axes. Note that pachytene is not reached yet at this stage, hence oocytes with fully formed synapsis are not represented in the graph. Counted cell numbers, standard deviation and weighted averages of percentages are shown from two experiments. Chi Square statistics indicates that ANKRD31 significantly alters the proportion of cells with distinct SYCP1 staining patterns. **** represents P<0.00001. (**C, D, F-H, L, N**) Indicated proteins were detected by immunofluorescence in nuclear surface spread spermatocytes (**C, D**) or oocytes (**F-H, L, N**). Early zygotene (**C**) or late pachytene (**D**) spermatocytes are shown. (**D**) Miniaturized H1t signal of the corresponding cell is shown in the bottom right corner of overlay images. Arrows mark two of the prominent flares of γH2AX on fully synapsed chromosome axes. Saturated γH2AX signal corresponds to the silenced chromatin of sex chromosomes (SB), which is a chromatin compartment where histone γH2AX hyper-accumulates in pachytene and diplotene stages. (**E**) Quantification of γH2AX flare numbers in late pachytene (late pa) and diplotene (diplo) spermatocytes. Three categories were distinguished, spermatocytes with less than 10 flares, between 10 and 40 flares and more than 40 flares of γH2AX on autosomes. Graph shows standard deviation and weighted averages of percentages of spermatocytes in the three categories from two experiments. Chi square statistics was used to calculate if loss of ANKRD31 significantly alters the proportion of cells with distinct numbers of γH2AX flares. (**F-H**) Zygotene oocytes are shown from 16 dpc fetuses. (**I-K**) Quantifications of DMC1 (**I**), RAD51 (**J**) and RPA (**K**) focus numbers in oocytes from 16 dpc foetuses. Counts are shown in early zygotene (e zy) and late zygotene cells that were split into two groups based on the condensation level of chromosome axes. Chromosome axis was fully formed, but axes were more condensed and more synapsed in the cells that were judged more advanced (l full axis) than in early stage cells with fully formed axes (e full axis). Number of counted cells and medians are indicated. Mann-Whitney U test calculated significance. (**L, N**) Late pachytene oocytes from newborn mice, where chromosome axes are fully synapsed, are shown. (**M**) Quantification of RPA focus numbers on late pachytene (lpa) and diplotene (di) oocytes of newborn mice of indicated genotypes. Counted number of cells and medians are shown. Mann-Whitney U test calculated significance. (**O**) Quantification of γH2AX staining patterns in late pachytene (late pa) and diplotene (diplo) oocytes. Four categories of γH2AX staining patterns were distinguished, oocytes with less than 10 flares, between 10 and 40 flares and more than 40 flares of γH2AX, and oocytes where intense γH2AX staining was detected throughout the nucleus (All over). Graph shows standard deviation and weighted averages of percentages of oocytes in the four categories from two experiments. Chi square statistics was used to calculate if loss of ANKRD31 significantly alters the proportion of cells with distinct numbers of γH2AX flares. (**C, D, F-H, L, N**) Bars, 10μm. (**A, B, E, I-K, M, O**) ns, *, ** and **** indicate no significance and significance at 0.05>P>0.01, 0.01 >P>0.001 and P<0.0001, respectively.

**Figure S6 related to Figure 6.**
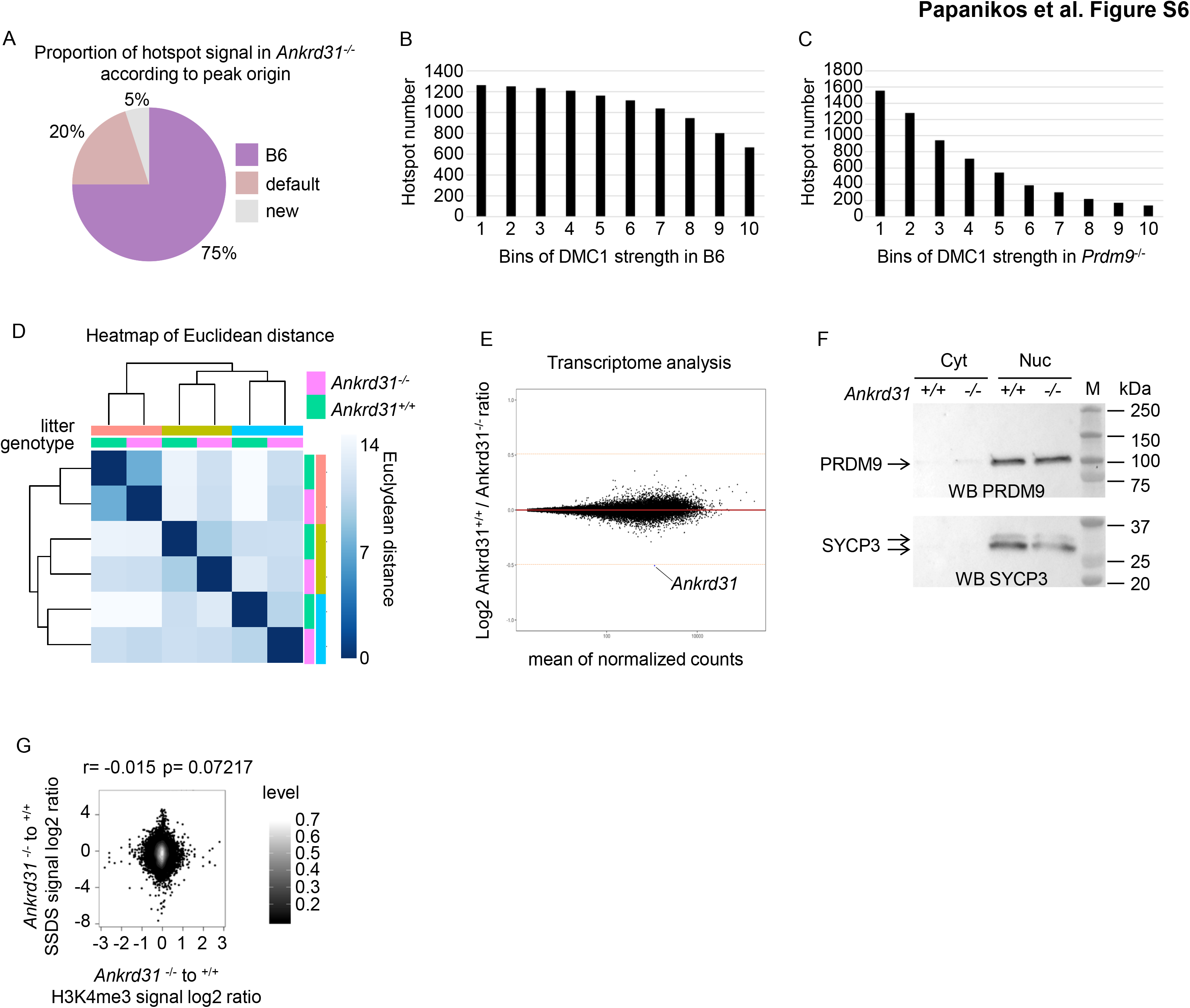
(**A**) Proportion of DMC1 ChIP SSDS signal from *Ankrd31^-/-^* hotspots that match PRDM9-defined (B6) or default hotspots, or that do not match either (new). (**B**) *Ankrd31^+/+^*-B6 hotspots were sorted into ten bins according to their strengths. Bin 1 and 10 represent the strongest and the weakest hotspots in *Ankrd31^+/+^* mice, respectively. Graph shows numbers of *Ankrd31^-/-^* hotspots that overlapped with *Ankrd31^+/+^*-B6 hotspots in each bin. (C) Default hotspots from *Prdm9^-/-^* were sorted into ten bins according to their strengths. Bin 1 and 10 represent the strongest and the weakest hotspots in *Prdm9^-/-^* mice, respectively. Graph shows numbers of *Ankrd31^-/-^* hotspots that overlapped with default hotspots of *Prdm9^-/-^* mice in each bin. (**D, E**) Transcriptomes of testes from 12 days old *Ankrd31^+/+^* and *^-/-^* mice were analyzed by RNA sequencing. Euclidean distance of testicular transcriptomes of three littermate *Ankrd31^+/+^* and *Ankrd31^-/-^* mice at 12 days of age. Heatmap indicates distance of transcriptomes. (E) Differential gene expression analysis (DESeq2) was used to calculate *Ankrd31^+/+^* to *^-/-^* expression ratios which are shown in log2 scale on the y-axis of the MA plot. The x-axis displays the mean expression. Lines indicate 1.4 fold change in expression. Blue color marks transcripts whose levels significantly change with at least 1.4 fold. Ankrd31 transcript is marked. (F) Nuclear (Nuc) and cytoplasmic (Cyt) extracts from testis of 12 days old mice were tested for the presence of PRDM9 and SYCP3 by western blot (WB). M: molecular weight marker. (G) Comparison of SSDS log2 signal change and H3K4me3 log2 signal change between *Ankrd31^-/-^* and *Ankrd31^+/+^* at each of PRDM9-defined hotspots. Spearman’s rank correlation coefficient (r) and associated p-value are shown. Levels represent the density calculated with the density2d function from R.

**Figure S7 related to Figure 7.**
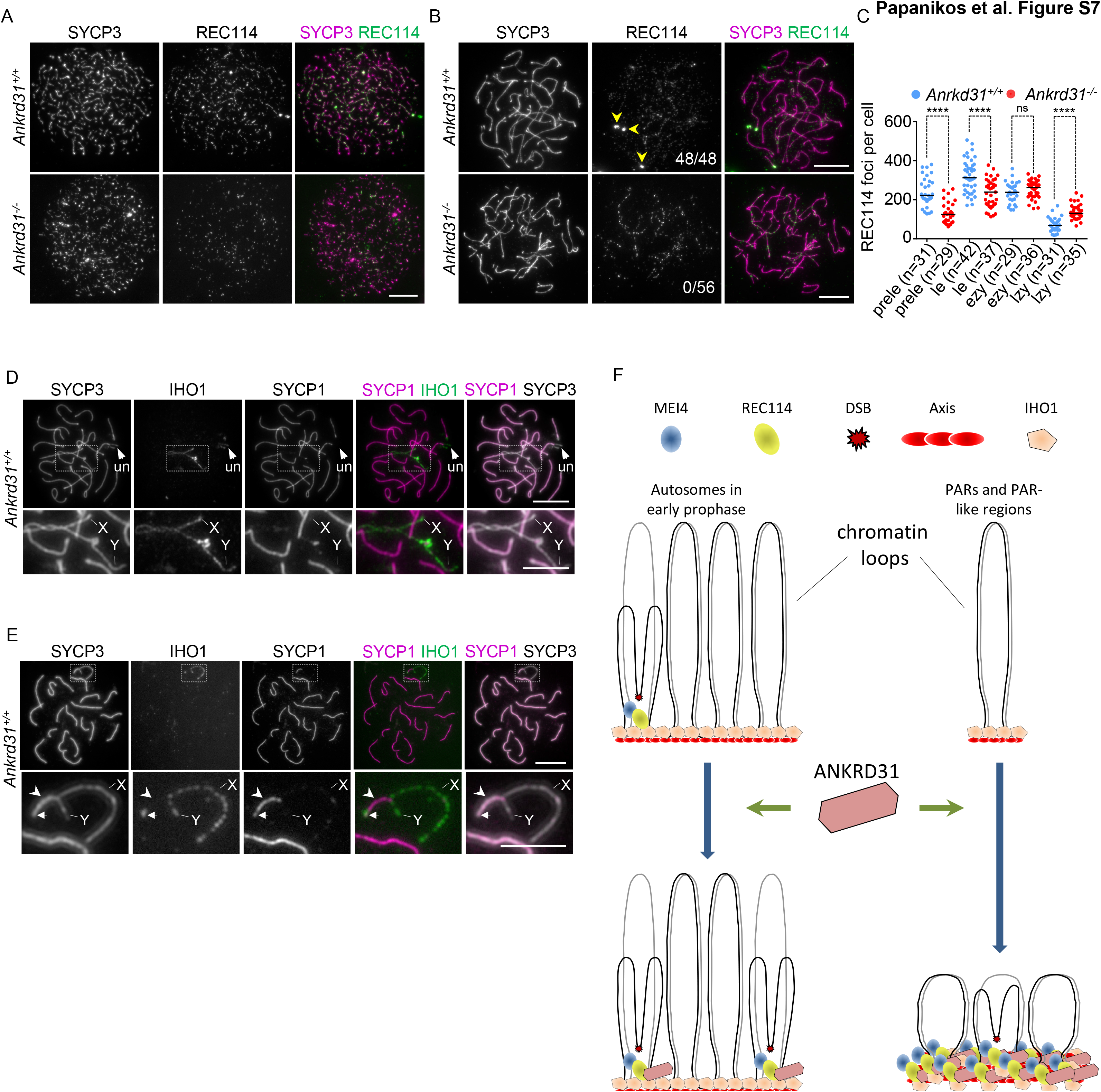
(**A, B, D, E**) Immunofluorescence was used to detect indicated proteins in nuclear spread spermatocytes of indicated genotypes. Leptotene, late zygotene, zygotene-to-pachytene transition, and early pachytene spermatocytes are shown in **A**, **B**, **D**, and **E**, respectively. (**A, B**) Matched exposure images of REC114 are shown. Arrowheads mark robust REC114 aggregates at the ends of three chromosomes. The fraction of late zygotene spermatocytes that contained REC114 aggregates is indicated in the REC114 image of **B**. (**C**) Quantification of REC114 focus numbers in preleptotene (prele), leptotene (le), early (ezy) and late (lzy) zygotene spermatocytes of indicated genotypes. Numbers (n) of examined cells and median focus numbers are indicated. Mann-Whitney U test calculated statistical significance, ns stands for no significance, **** indicates significance at P<0.0001. (**D, E**) Insets in the images of full nuclei (top panels) are shown at higher magnification in bottom panels. Axes of chromosome X and Y are marked. An unsynapsed region in an autosome is pointed out (un) in **D**. (**E**) Arrow points at the site of persisting IHO1 signal on the synapsed PAR-end of sex chromosomes, and arrowhead marks the extended region of synapsis that probably connects non-homologous sections of X and Y chromosomes. (**F**) Graphical representation of key ANKRD31 functions within the previously proposed loop-axis model of DSB formation (Blat et al., 2002; Panizza et al., 2011). Blue block arrows represent transition between states of chromatin and the axis, green arrows represents stimulation by ANKRD31 Black and grey lines represent DNA of chromatin loops of sister chromatids that are tethered to axes. The extent of alignment of sister DNA sequences in loops is uncertain. It is also uncertain whether either both or one sister chromatid loop is recruited to the DNA break promoting complex on axes.These uncertainties are not pertinent to our understanding of ANKRD31 functions hence we represent only one of the possible scenarios. (Left) Outside of PAR or PAR-like regions DSB-promoting proteins including MEI4 and REC114 form pre-DSB recombinosomes tethered to axes by a spread out platform of axis-bound IHO1. DSB formation involves the recruitment of loop sequences to axis-bound pre-DSB recombinosomes. ANKRD31 associates with the IHO1-dependent pre-DSB recombinosomes and enhances their formation. Thereby, ANKRD31 permits timely formation of well-distributed DSB-breaks, which improves efficiency of homolog pairing and SC formation. (Right) In the PAR, ANKRD31 permits the assembly of a disproportionately long (Acquaviva, Jasin & Keeney, personal communications) and robust axis where DSB-promoting proteins are recruited with high efficiency in an IHO1-independent manner. The resultant short DNA loops and high concentration of DSB-promoting proteins on the axis ensure the formation of obligate DSBs in the PAR.

**Table S1 is related to Figure 1.**
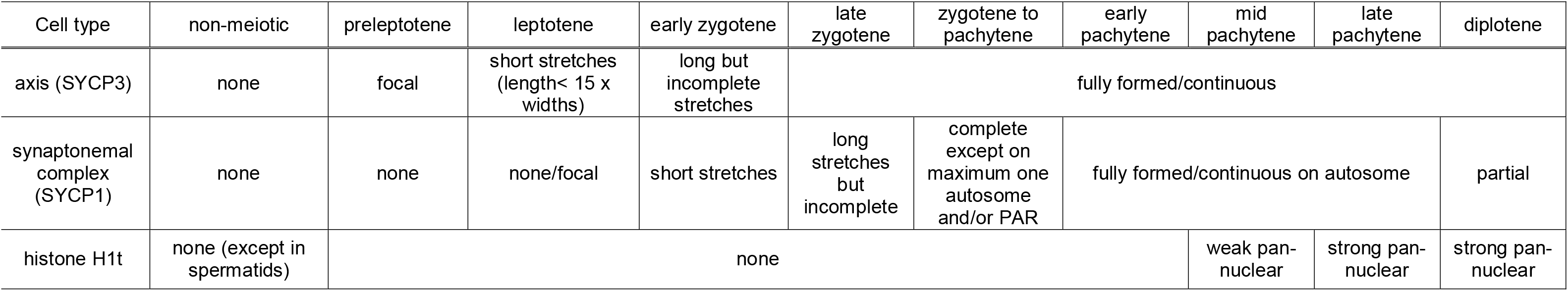
A summary of distinguishing features of meiotic prophase substages. First row lists substages that can be distinguished by the combining detection of axis (SYCP3, second row), synaptonemal complex (SYCP1, second row) and histone H1t (fourth row).

**Table S2 is related to Figure 2.** Summary of yeast two-hybrid (Y2H) interactions between fragments of ANKRD31 and indicated proteins. The first column indicates if ANKRD31 fragments were tested for interaction in a Y2H bait or prey vector. Second column indicates positions of N- and C-terminal amino acids of ANKRD31 fragments along the full length ANKRD31 protein. Third column indicates the presence or absence of conserved domains of ANKRD31 in tested fragments. Interactions were tested by growing yeast that carry the appropriate bait and prey vectors at 30L C on selective plates. Growth was inspected 2 and 3 days after plating droplets of 0.5 OD budding yeast cell suspensions on selective plates. The scoring of growth is shown in the last four columns, - represents no growth after three days, + growth of a lawn or multiple colonies after three days, ++ growth of a lawn after two days, NA indicates not tested interaction.

| ANKRD31 |  |  | Interactions tested with |  |  |  |
| --- | --- | --- | --- | --- | --- | --- |
| GAL4-BD/GAL4-AD | Position (aa) | Motif | REC114 | IHO1 | MEI4 | HORMAD1 |
| -/+ | 1-473 | partial Ankyrin repeat 1 | - | + | - | - |
| -/+ | 1-375 | no motif | - | - | NA | NA |
| -/+ | 376-473 | partial Ankyrin repeat 1 | - | - | NA | NA |
| -/+ | 376-821 | Ankyrin repeat 1 | + | ++ | - | - |
| -/+ | 460-606 | Ankyrin repeat 1 | + | + | NA | NA |
| -/+ | 570-821 | partial Ankyrin repeat 1 | - | - | - | - |
| -/+ | 750-1150 | no motif | - | - | - | - |
| -/+ | 1055-1459 | Ankyrin repeat 2, coiled coil domain | - | - | - | - |
| +/+ | 1262-1857 | coiled coil domain, CR4, CR5 | ++/++ | NA/- | NA/- | NA/- |
| +/+ | 1260-1460 | coiled coil domain | -/- | NA |  |  |
| +/+ | 1460-1790 | CR4 | -/- |  |  |  |
| +/+ | 1784-1857 | CR5 | +/+ |  |  |  |

**Table S3 is related to Figure 2.**
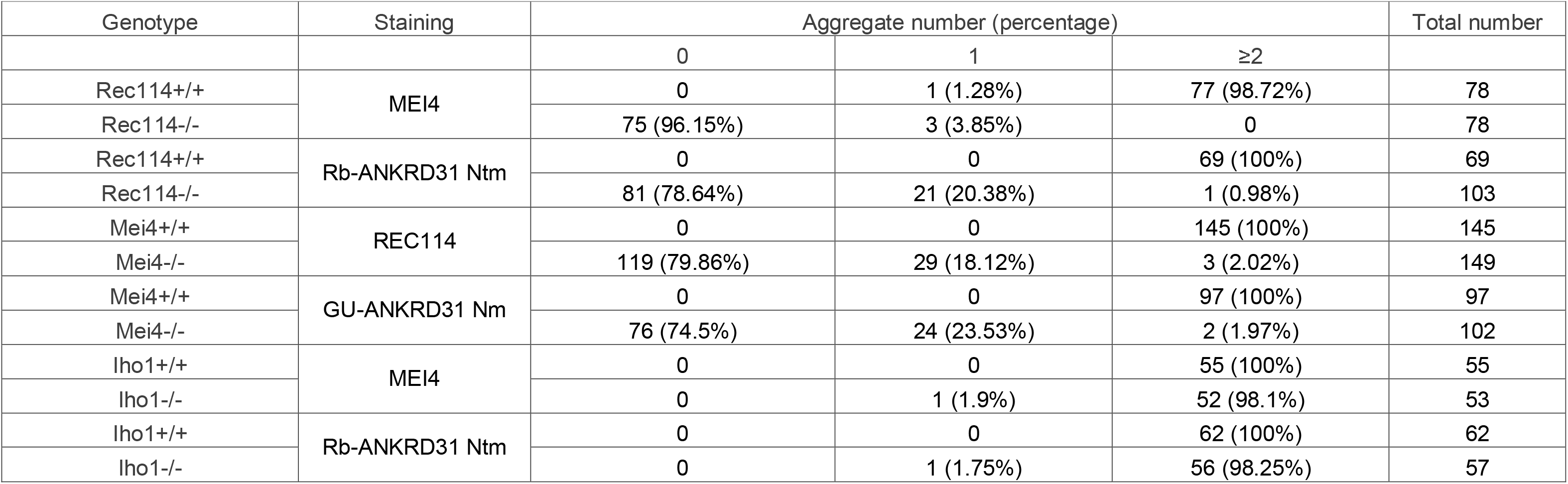
Quantification of aggregates of indicated proteins (column “staining”) in zygotene-like spermatocytes of the indicated genotype (first column). Number and percentage (in brackets) of spermatocytes that had no aggregates, had one aggregate, or had at least two aggregates associated with ends of chromosome axes are shown in columns 3-4. Last column shows total number of observed spermatocytes.

**Table S4 is related to Figure 3.**
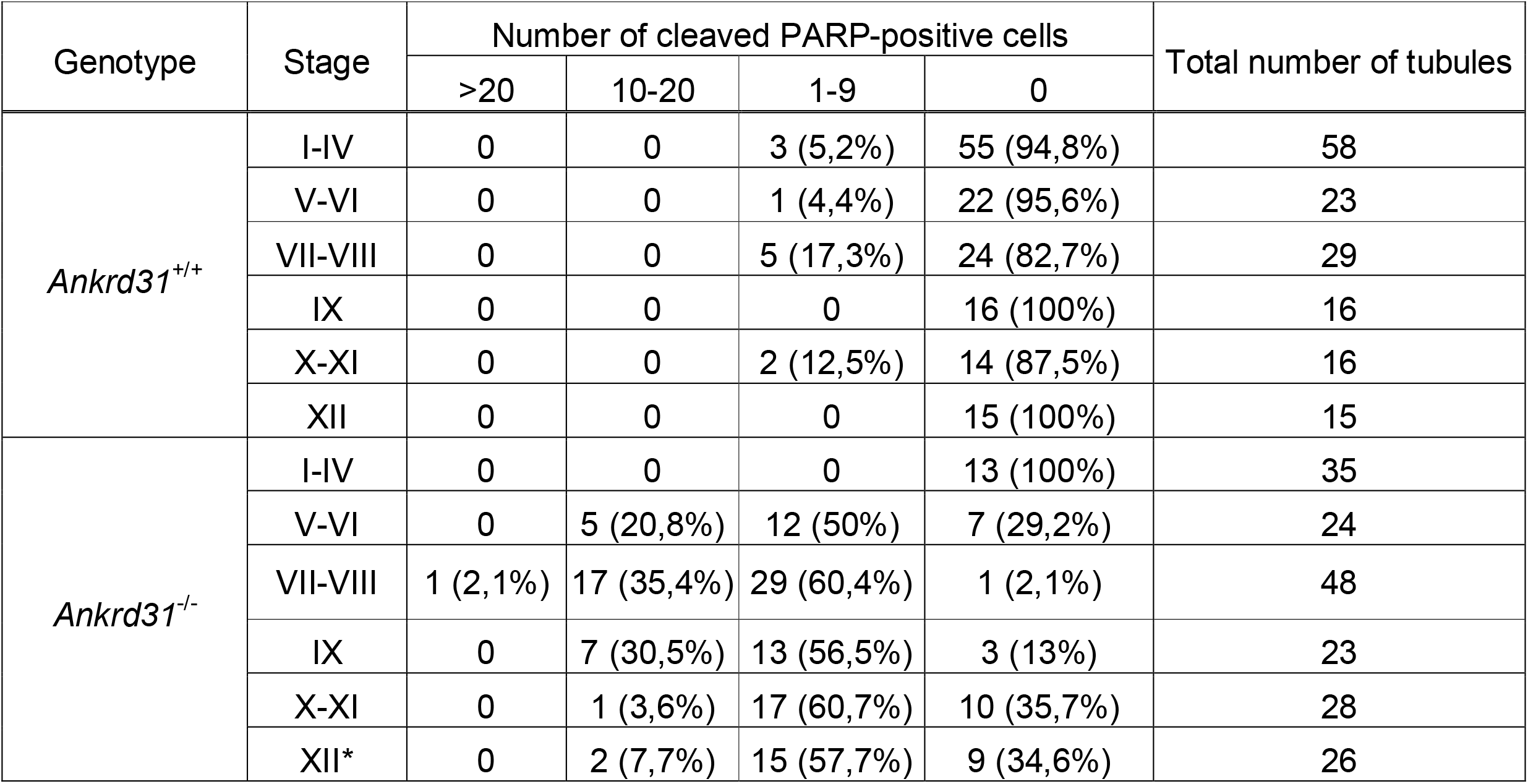
Quantification of cleaved PARP positive cells in luminal layers of spermatocytes in seminiferous tubules of adult *Ankrd31^+/+^* and *Ankrd31^-/-^* mice. Quantification was based on chromatin morphology of spermatogenic cells as detected by DAPI in cryosections of testes. Cleaved PARP and histone H1t were detected by immunofluorescence. Histone H1t is expressed in spermatocytes from mid pachytene, hence histone H1t staining facilitated easy identification of stage V-XII tubules, which contain spermatocytes beyond mid pachytene. Columns 3-6 show the numbers and the percentages (in brackets) of seminiferous tubules that had more than 20, 10-20, 1-9 or no cleaved PARP positive spermatocytes. The last column indicates the total number of inspected seminiferous tubules from testes of two mice of each genotype.

**Table S5 is related to Figure 3.** Quantification of synapsis in oocytes of 18 dpc *Ankrd31^+/+^* and *Ankrd31^-/-^* fetuses. Table shows the number and the percentage (in brackets) of oocytes where the indicated numbers of chromosomes (in leftmost column) are fully synapsed (first column for each genotype), partially (second column for each genotype) or fully unsynapsed (third column for each genotype). Note that percentages from different columns do not add up to 100% for each genotype because some cells are counted twice due to the presence of both partially and fully unsynapsed chromosomes in the same cell.

|  | *Ankrd31*<sup>+/+</sup> |  |  | *Ankrd31*<sup>-/-</sup> |  |  |
| --- | --- | --- | --- | --- | --- | --- |
| Number of chr. | Fully synapsed | Partially unsynapsed | Fully unsynapsed | Fully synapsed | Partially unsynapsed | Fully unsynapsed |
| 0 | 0 | 105 (78.35%) | 128 (95.5%) | 2 (1.74%) | 62 (53.92%) | 77 (66.95%) |
| 1 | 0 | 16 (11.93%) | 3 (2.25%) | 0 | 13 (11.31%) | 15 (13.04%) |
| 2 | 0 | 7 (5.22%) | 2 (1.5%) | 0 | 19 (16.52%) | 10 (8.7%) |
| 3 | 0 | 3 (2.25%) | 1 (0.75%) | 2 (1.74%) | 7 (6.08%) | 2 (1.74%) |
| 4 | 0 | 2 (1.5%) | 0 | 1 (0.87%) | 4 (3.48%) | 4 (3.48%) |
| 5 | 0 | 0 | 0 | 0 | 3 (2.61%) | 0 |
| 6 | 0 | 1 (0.75%) | 0 | 1 (0.87%) | 5 (4.34%) | 1 (0.87%) |
| 7 | 0 | 0 | 0 | 1 (0.87%) | 1 (0.87%) | 1 (0.87%) |
| 8 | 0 | 0 | 0 | 0 | 1 (0.87%) | 1 (0.87%) |
| 9 | 0 | 0 | 0 | 1 (0.87%) | 0 | 0 |
| 10 | 0 | 0 | 0 | 2 (1.74%) | 0 | 0 |
| 11 | 0 | 0 | 0 | 0 | 0 | 0 |
| 12 | 0 | 0 | 0 | 1 (0.87%) | 0 | 0 |
| 13 | 0 | 0 | 0 | 0 | 0 | 0 |
| 14 | 1 (0.75%) | 0 | 0 | 5 (4.34%) | 0 | 2 (1.74%) |
| 15 | 2 (1.5%) | 0 | 0 | 6 (5.22%) | 0 | 0 |
| 16 | 1 (0.75%) | 0 | 0 | 7 (6.08%) | 0 | 1 (0.87%) |
| 17 | 4 (3%) | 0 | 0 | 6 (5.22%) | 0 | 0 |
| 18 | 8 (6%) | 0 | 0 | 15 (13.04%) | 0 | 1 (0.87%) |
| 19 | 14 (10.4%) | 0 | 0 | 9 (7.83%) | 0 | 0 |
| 20 | 104 (77.6%) | 0 | 0 | 56 (48.7%) | 0 | 0 |
| total number of cells | 134 |  |  | 115 |  |  |

**Table S6 is related to Figure 6.** Summary of DMC1 ChIP-seq and histone H3K4me3 ChIP-seq experimental design and associated mapping results.

| Genotype | Antibody | Experiment | Replicate | Sequencing | Mapped reads * |
| --- | --- | --- | --- | --- | --- |
| B6 | H3K4me3 | ChIPseq | 1 | SE75bp | 94,782,757 |
| B6 | H3K4me3 | ChIPseq | 2 | SE75bp | 106,004,616 |
| B6 | H3K4me3 | Input | 1 | SE75bp | 123,846,185 |
| B6 | H3K4me3 | Input | 2 | SE75bp | 134,577,035 |
| Ankrd31<sup>-/-</sup> | H3K4me3 | ChIPseq | 1 | SE75bp | 100,025,923 |
| Ankrd31<sup>-/-</sup> | H3K4me3 | ChIPseq | 2 | SE75bp | 102,192,500 |
| Ankrd31<sup>-/-</sup> | H3K4me3 | Input | 1 | SE75bp | 130,190,990 |
| Ankrd31<sup>-/-</sup> | H3K4me3 | Input | 2 | SE75bp | 132,645,159 |
| B6 | DMC1 | SSDS | 1 | PE50bp | 7,879,020 |
| B6 | DMC1 | SSDS | 2 | PE50bp | 17,221,289 |
| Ankrd31<sup>-/-</sup> | DMC1 | SSDS | 1 | PE50bp | 17,129,611 |
| Ankrd31<sup>-/-</sup> | DMC1 | SSDS | 2 | PE50bp | 6,537,666 |

\* Uniquely, deduplicated, high quality (>30) mapped reads (only single-stranded DNA for SSDS experiments).
SE, single-end. PE, Paired-end. SSDS, Single-Strand DNA Sequencing. ChIPseq, Chromatin Immunoprecipitation and sequencing.

**Table S7 is related to Figure 6.** Quantification of DMC1 ChIP SSDS and histone H3K4me3 ChIP-seq signal (reads per million per kilobase, rpkm), in the non-PAR part of X chromosome, the PAR and two PRDM9-independent hotspots in the vicinity of the PAR.

| Region | Coordinates | DMC1 |  |  | H3K4me3 |  |  |
| --- | --- | --- | --- | --- | --- | --- | --- |
|  |  | *Ankrd31*<sup>+/+</sup> | *Ankrd31*<sup>-/-</sup> | ratio +/+ over -/- | *Ankrd31*<sup>+/+</sup> | *Ankrd31*<sup>-/-</sup> | ratio +/+ over -/- |
| nonPAR chrX | chrX:0-169969759 | 0.151 | 0.157 | 0.963 | 0.114 | 0.121 | 0.940 |
| PAR X | chrX:169969759-170931299 | 0.608 | 0.0644 | 9.441 | 0.129 | 0.173 | 0.747 |
| 1st hotspot upstream of PAR | chrX:169926497-169928227 | 15.535 | 2.009 | 7.734 | 1.943 | 2.274 | 0.855 |
| 2nd hotspot upstream of PAR | chrX:169878711-169881527 | 13.208 | 1.987 | 6.648 | 4.780 | 5.443 | 0.878 |

